# The role of arbuscular mycorrhiza and organosulfur mobilizing bacteria in plant sulphur supply

**DOI:** 10.1101/2021.02.08.429889

**Authors:** Jacinta Gahan, Orla O’Sullivan, Paul Cotter, Achim Schmalenberger

## Abstract

AM fungi are enhancing growth and health of many land plants but only some of these beneficial mechanisms are well understood. This study aimed to uncover the role of bacteria colonising AM fungi in organically-bound sulfur (S) mobilisation, the dominant S pools in soil that are not directly available to plants. The effect of an intact AM symbiosis with access to stable isotope organo-^34^S enriched soils encased in 35 µm mesh cores was tested in microcosms with *Agrostis stolonifera* and *Plantago lanceolata*. At 3 month intervals, the plant shoots were analysed for ^34^S uptake. After 9 months, hyphae and associated soil was picked from static (mycorrhizal) and rotating (severed hyphae) mesh cores and corresponding rhizosphere soil was sampled for bacterial analysis. AM symbiosis increased uptake of ^34^S from organo-^34^S enriched soil at early stages of plant growth when S demand appeared to be high. The static (mycorrhizal) treatments were shown to harbour larger populations of cultivable heterotrophs and sulfonate mobilising bacteria. Microbial communities were significantly different in the hyphosphere of mycorrhizal hyphae and hyphae not associated to plant hosts. Sulfate ester (arylsulfatase enzyme assay, *atsA* gene) and sulfonate mobilising activity (*asfA* gene) was altered by an intact AM symbiotic partnership which stimulated the genera *Azospirillum, Burkholderia* and *Polaromonas*. Illumina sequencing revealed that AM symbiosis led to community shifts, reduced diversity and dominance of the Planctomycetes and Proteobacteria. This study demonstrated that AM symbioses can promote organo-S mobilization and plant uptake through interaction with hyphospheric bacteria.

**Research highlights:**

- AM hyphae enhanced uptake of organically bound ^34^S at early stages of growth.
- AM hyphosphere harboured a large population of organo-S desulfurizing bacteria.
- Microbial communities significantly differed in rotating and static mesh cores.
- AM hyphae influenced bacterial sulfate ester and sulfonate mobilising activity.
- AM hyphae reduced bacterial diversity, increased Planctomycetes and Proteobacteria abundance.

## 1. Introduction

The essential macro-nutrient sulfur (S) is increasingly limiting to crop yield and quality as a result of reduced atmospheric deposition while high yielding crop varieties rapidly deplete soil S stocks. These factors are exacerbated by the fact that S present in soil is approximately 95% organically bound as sulfate-esters and sulfonates (Autry & Fitzgerald, 1990, Kertesz & Mirleau, 2004) and not directly available to plants which rely upon soil microbial populations for organo-S mineralisation (Kertesz *et al*., 2007). The ability to mineralise sulfate esters is common for many bacteria and saprotrophic fungi in soil and the *atsA* gene is one such marker for this ability (Beil *et al*., 1995, Klose *et al*., 1999). A bacterial multicomponent mono-oxygenase enzyme complex was discovered that is capable of mobilising a large variety of sulfonates of which the *asfA* and *ssuD* genes are markers for aromatic and aliphatic sulfonate mobilisation, respectively (Vermeij *et al*., 1999, Kertesz & Mirleau, 2004). The *atsA* gene for sulfate ester mineralisation has been most comprehensively studied for *Pseudomonas* species (Kahnert *et al*., 2000, Kahnert *et al*., 2002). The sulfonate mobilising *asfA* gene has been identified in the Beta-Proteobacteria; *Variovorax, Polaromonas, Hydrogenophaga, Cupriavidus, Burkholderia* and *Acidovorax*, the Actinobacteria; *Rhodococcus* and *Williamsia*, and the Gamma-Proteobacteria; *Pseudomonas* and *Stenotrophomonas* (Schmalenberger & Kertesz, 2007, Schmalenberger *et al*., 2008, Schmalenberger *et al*., 2009, Fox *et al*., 2014, Gahan & Schmalenberger, 2015).

Soil S cycling involves complex interactions between free living and symbiotic rhizospheric microbial populations. AM fungi are one such microbial population that form an endosymbiosis with 77% of angiosperms, 45% of 84 species of gymnosperms and 52% of 400 species of fern and lycopod (Wang & Qiu, 2006). Their characteristic structure, the arbuscule, is extensively branched and acts as an efficient site for metabolite exchange with the host plant (Smith & Read, 1997). Additionally, their extraradical hyphae (ERH) play an important role in nutrient acquisition via extensive networks of microscopic hyphae that access a volume of soil orders of magnitude greater than plant roots (Nagahashi & Douds, 2000). Studies have shown that the presence of AM fungi enhances S uptake for maize, clover (Gray & Gerdemann, 1973) and tomato (Cavagnaro *et al*., 2006). More recently, the presence of the AM fungus *Rhizophagus irregularis* (formerly known as *Glomus intraradices*) on transformed carrot roots demonstrated uptake of reduced forms of S *in vitro* (Allen & Shachar-Hill, 2009). Rates of uptake and transfer of reduced S were comparable to that of SO_4_^2-^ when the latter was limited. Uptake of SO_4_^2-^ in the rhizosphere leads to a zone of depletion analogous to that observed for phosphorus (Buchner *et al*., 2004). The AM fungal ERH could extend out past this zone of SO_4_^2-^ depletion and play an important role in provision of S under conditions of limitation (Kertesz *et al*., 2007). Additionally, recent investigations have revealed that AM fungi influence the expression of plant sulfate transporters and as a consequence improve S uptake (Giovannetti *et al*., 2014).

AM symbiosis alters the rhizospheric microbial communities composition by modifying the biochemical composition of root exudates (Barea *et al*., 2002, Boer *et al*., 2005). Additionally, translocation of energy rich C compounds to the extended soil environment via their extensive ERH networks provides an important niche for functional interactions (Gryndler *et al*., 2000). Diverse soil microbial communities are essential for soil fertility and plant vitality (Gianinazzi & Schüepp, 1994, Siciliano *et al*., 2014) and AM hyphae in a native grassland ecosystem have been shown to host a larger community of sulfonate mobilising bacteria with potential to improve plant S supply (Gahan & Schmalenberger, 2015). Moreover, it has been shown that the addition of morpholine ethane sulfonic acid to soil stimulated sulfonate mobilising bacteria and their metabolites enhanced ERH growth of *R. irregularis* (then *G. intraradices* (Vilarino *et al*., 1997)). This is important for maximising S uptake as enhanced hyphal growth stemming from sulfonate mobilising bacterial metabolites may further stimulate the proliferation of this community in a potential positive feedback loop.

The hypothesis of this study was that the presence of an intact AM symbiosis would increase uptake of S from organo-^34^S enriched soil. The mechanism for which lies in interactions with organo-S mobilising microbes that increase organo-S mineralization for enhanced uptake of this newly mineralised S through AM hyphae. This hypothesized activity is important to overcome plant S limitations and to reduce over-dependencies on synthetic fertilizer use.

## 2. Materials and Methods

### 2.1 Site description

The soil used to create the experiments was obtained from Teagasc, Johnstown Castle, Wexford, Ireland (52°16’N, 6°30’W, 30 m above sea level). The soil type is a poorly drained gley soil (pH 6), organic matter (11%), loamy topsoil (18% clay) and classified as Mollic Histic Stagnosol (WRB 2006). The soil has not received P or S fertiliser since 1968 and has not been ploughed since 1970 (P0-0, site 5A). Swards are mixtures of *L. perenne, D. glomerata* and various meadow grass species (Tunney *et al*., 2010).

#### 2.2. Stable isotope enriched soil

The most common form of elemental S in nature is ^32^S (95.02%) and stable isotope ^34^S represents 4.21% of S existing naturally in soil. For this study, S uptake and incorporation into plant biomass was undertaken using the ratio of ^34^S to ^32^S (δ^34^S values) (Krouse *et al*., 1996).

To enrich the soil with organo-^34^S, 20 g soil aliquots were treated with ^34^S-SO_4_ (20 mg/kg soil) and glucose (6 g/kg) to stimulate S immobilisation and optimise the amount of S in the C-bonded fraction (Dedourge *et al*., 2004). The soil was incubated for 6 weeks at 25 °C and aerated fortnightly via aggregation. After 6 weeks, soil was thoroughly mixed with 20 ml of 0.01 M CaCl_2_ for 30 min on a Elmi Intelli-Mixer RM-2 (Elmi Tech Ltd, Latvia) and subjected to centrifugation in an Allegra X-22R centrifuge (Beckman Coulter, USA) at 3900 rcf for 20 min and to remove inorganic S (Dedourge *et al*., 2004). This process was repeated two additional times before the supernatant was discarded and the residual material was dried, ground and brought to 60% water holding capacity (WHC). To counteract the loss of essential nutrients, the enriched fraction was supplemented with Hoagland’s solution (S free, 50%) (Hoagland & Snyder, 1933). The ^34^S enriched soil samples were transferred into nylon mesh cores (up to 6 g per core) with windows of either 35 µm or 1 µm (Johnson et al. 2001).

For a second experiment, sulfonate enriched organo-^34^S was used and this was prepared as above with the addition that the ^34^S-enriched soil was treated with a sulfatase enzyme from *Helix pomatia* (10 U g^-1^ soil; Sigma Aldrich, St Louis, MO, USA) and mixed on an Elmi Intelli-Mixer RM-2 (10 rpm; Elmi Tech Ltd, Latvia) at 37 °C for 4 h. After which, the soil was treated with 0.01 M CaCl_2_ and centrifuged as described above. This was repeated two additional times before the residual material was dried and adjusted to 60 % WHC. Soil can contain up to 60% of its S as sulfate esters (Autry and Fitzgerald 1990) and the quantity of sulfatase enzyme used was calculated on this basis. Hoagland’s solution (50%, S free) was used to counteract nutrient losses other than S.

### 2.3. Core design and constructing the soil microcosms

Cores were constructed using acrylonitrile butadiene styrene (ABS) water pipe (150 mm height, 18 mm diameter). Two windows were cut into each core, the size of which constituted 90% of the surface area of the core (Supplementary Figure S1). Nylon mesh (Plastok Associates Ltd, UK) with 35 µm pores was used to cover the windows of the core and 1 µm nylon mesh was used to cover the base. The mesh was attached using Tensol No. 12 adhesive (Evode Speciality Adhesives Ltd, UK) and the completed cores were cured (24 h, 80 °C). The 35 µm mesh is sufficiently large to allow mycorrhizal colonisation while preventing root access. A permeable base of 1 µm allows free drainage of water through the cores. This set of cores was used to analyse the role of AM in uptake of organo-^34^S and sulfonates^34^S specifically. Additional cores with 1 µm mesh windows were created to determine the effect of leaching from the cores.

#### 2.3.1. Organo-^34^S enriched soil microcosms

Microcosm systems were constructed from cuttings of Plexiglas (12 x 11 x 2.5 cm, chemically welded with Ethylene Dichloride) and filled with 230 g of a sand (Glenview Natural Stone, Ireland) and soil (Teagasc, Johnstown Castle) mixture (one part sand, one part soil w/w). These were planted with *A. stolonifera* and *P. lanceolata* respectively and inoculated with 2 g of *Rhizophagus irregularis* (purchased as *Glomus intraradices;* Symbivit, Symbion, Czech Republic). For this experiment, two mesh cores, containing 6 g of organo-^34^S stable isotope enriched soil, were inserted into each soil microcosm (Supplementary Figure S2). The organo-^34^S enriched cores were either rotated twice weekly to sever AM hyphae (rotating) or undisturbed (static) to allow AM colonisation. Each treatment was carried out in replicates of six. The microcosms were grown in an A1000 Adaptis plant growth chamber (Conviron, Germany) with day-night temperature of 25-15 °C, respectively, 12 h day, 70% RH, and 320 µmoles m^-2^ s^-2^ PAR. The systems were watered with dH_2_O three times a week and once fortnightly supplemented with half times S free Hoagland’s solution (Hoagland and Snyder 1933).

#### 2.3.2. Sulfatase treated organo-^34^S enriched soil microcosms

Soil microcosms with *A. stolonifera* were established as above but cores for this experiment contained sulfatase treated organo-^34^S enriched soil. The soil was inserted into 35 µm mesh cores in *A. stolonifera* microcosms (6 static, 6 rotating). These systems were established to determine the AM role in mobilisation of ^34^S-sulfonates in the absence of sulfate esters. The systems were grown under the same conditions as described above.

#### 2.3.3. Organo-^34^S enriched soil microcosms with 1 µm mesh cores

The ^34^S stable isotope enriched soil was prepared as outlined above, but the core windows were covered with 1 µm mesh which is impenetrable by AM hyphae. This experiment was undertaken to quantify the effect of leaching from the organo-^34^S enriched mesh cores. Replicates of 6 soil microcosms with *A. stolonifera* as host plant were established and the 1 µm mesh cores were inserted as previously outlined and were grown under controlled conditions as described above.

### 2.4. Determination of ^34^S uptake

Determination of ^34^S uptake was achieved using Elemental Analysis -Isotope Ratio Mass Spectrometry (EA-IRMS) undertaken by Iso-Analytical (Chesire, UK) as outlined in the supplementary information section S1. At 3 month intervals, the aboveground biomass was destructively harvested to analyse the S flow from labelled organo-^34^S in the mesh cores to its incorporation into plant tissue. The plant biomass was dried at 80 °C for 5 d, ground to a fine powder and a 0.5 g aliquot was subjected for EA-IRMS. This was repeated at 6 and 9 months at which point the systems were deconstructed and subjected to downstream cultivation dependent and independent experimentation. Additionally, at 9 months one of the two cores in each *A. stolonifera* microcosm was taken and re-planted into new system to ascertain long term organo-^34^S uptake. EA-IRMS of aboveground biomass was undertaken at 3 months for the sulfatase treated and 1 µm mesh core microcosms.

### 2.5. S K-edge X-ray absorption near edge spectroscopy

XANES analysis was used to identify the chemical oxidation states of S, reduced (sulfides, thiols), intermediate (sulfoxides, sulfonates) and oxidised (sulfate esters, sulfates) as described previously (Schmalenberger *et al*., 2011). In the present case, soil subsamples from two static and two rotating core experiments of *A. stolonifera* were compared. Soil samples were dried at 105 °C for 12 h and were ground to a fine powder using a pestle and mortar. SO_4_ ^2-^ was extracted from each of the samples with 0.01 M CaCl_2_ using the same approach highlighted above. The soil was freeze dried in a Labconco FreeZone™ 4.5 L Freeze dry system (Labconco, MO, USA). XANES was carried out on the KMC-1 beam line at BESSY II (Helmholtz-Centre Berlin for Materials and Energy, Berlin, Germany, for further details see supplementary information S2). The KMC-1 is a soft x-ray double crystal monochromator beamline for the energy range 2-12 keV.

### 2.6. Percentage root colonisation

At the time of harvest, each plant and treatment was examined for root colonisation by AM fungi using a modified version of the grid line intersect method (McGonigle *et al*., 1990) as described in the supplementary information S3. Arbuscular colonization (AC) and vesicular colonization VC, respectively, were calculated by dividing their respective counts by the total number of intersections examined. Hyphal colonization (HC) was calculated as a proportion of the positive intersections. For each organo-^34^S stable isotope soil microcosm, 3 replicates of 100 fields of view per treatment, per plant were examined to calculate AM fungal colonisation.

### 2.7. Extraction and quantification of bacteria from AM hyphae

To harvest the microcosms, 1 g of hyphae with adhering soil (hyphosphere) was taken from each core and 1 g of roots with adhering soil (rhizosphere) was analysed in parallel. The hyphosphere and rhizosphere bacteria were extracted into 10 mL of sterile saline solution (0.85%) and rotated at 75 rpm on an Elmi Intelli-Mixer RM-2 (Elmi Tech Ltd, Latvia) (30 min, 4 °C). A 0.1 ml aliquot of the resulting suspension was used for bacterial community quantification (cultivation dependent). The remainder of the suspensions were centrifuged at 4500 rpm (4 °C, 20 min) and the obtained pellets were immediately frozen (−18 °C) for subsequent cultivation independent analysis (see below).

The aliquot of bacterial suspension from above was used for tenfold serial dilutions in saline (0.85% w/v). MPN analysis was undertaken in agar-free R2A (Reasoner *et al*., 1979), MM2TS and MM2LS ((Fox *et al*., 2014); supplementary information S4)) to enumerate the cultivable heterotrophic, sulfonate and polymeric sulfonate mobilising communities, respectively. A 20 µL aliquot from each dilution (10^1^-10^7^) was added to 200 µL of either R2A MM2TS or MM2LS in 96 well microtitre plates. Following 2 weeks of growth in an Innova Incubator Shaker Series (New Brunwick Scientific, UK) (75 rpm, 25 °C), the OD_590_ was recorded after 3 min shaking (Intensity Level 3) using an ELX808IU spectrophotometer (Bio Tek Instruments Inc., Winfrom two static and two rotating core experiments of A. stolonifera were compared. Soil ooski, VT). The OD_590_ was used to identify the lowest dilution with growth in all five wells, the number of wells with growth in the subsequent two dilutions was used to generate a three digit number and this was used to obtain an MPN g^-1^ value (FDA, 2011). This MPN g^-1^ value was substituted into the following equation: MPN g^-1^ x (1/V) x (DF). V is the volume in mL inserted into each well and DF is the dilution factor (0.02 mL (V) and 20 (DF) for this experiment).

### 2.8. Community fingerprinting

Community fingerprinting analysis was undertaken for the organo-^34^S microcosms. The frozen pellets of the bacterial extractions were used for community DNA extractions using the UltraClean Soil DNA extraction kit from MoBio as described by the manufacturer (Carlsbad, CA). Subsequent DGGE analysis was carried out for the bacterial 16S rRNA, 18S AM fungal and ITS saprotrophic fungal communities. PCR and DGGE conditions as well as a list of the selected primers for PCR-DGGE are available in the supplementary information S5 and supplementary table S1.

### 2.9. Arylsulfatase activity

A calorimetric method of assaying soil arylsulfatase activity was used to quantify arylsulfate ester cleaving enzymes in soil (Tabatabai & Bremner, 1970). Acetate buffer (0.5 M, pH 5.8) was prepared by dissolving 64 g of sodium acetate trihydrate in 200 mL dH_2_O, adding 1.70 mL glacial acetic acid (99%) and diluting this to 1 L. *p*-Nitrophenol solution (500 µg/L) was prepared in acetate buffer.

Arylsulfatase activity was analysed for hyphosphere and rhizosphere, static and rotating treatments. For analysis, 1 g of sieved (2 mm) soil was placed in 15 mL centrifuge tubes with 4 mL of acetate buffer, 0.25 mL of toluene and 1 mL of 20 mM *p*-nitrophenyl sulfate. The contents were vigorously mixed and incubated at 37 °C for 1 h (rotated 360 degrees at 10 minute intervals). After the incubation, 2 mL of 1 M NaOH and 1 mL of 0.5 M CaCl_2_ were added to stop the reaction. The centrifuge tubes were subjected to centrifugation (10 min, 4500 rpm). The absorbance of the supernatant was recorded at 400 nm in a UV MINI spectrophotometer (Shimadzu, Japan) (Tabatabai & Bremner, 1970).

The *p*-Nitrophenol content of the treatments was determined in reference to a calibration curve. This curve was obtained by preparing 100 mL of *p*-Nitrophenol solution (500 µg/L), of this 0, 1, 2, 3, 4, and 5 mL aliquots were prepared and made up to 5 mL in H_2_O to generate the 0, 10, 20, 30, 40, and 50 µg *p*-Nitrophenol standards. At this point, the standards were treated as described above for the post incubated soil samples and the absorbance of the supernatant was recorded at 400 nm in a UV MINI spectrophotometer (Shimadzu, Japan).

### 2.10. Diversity of asfA and atsA gene

Diversity of the sulfonate mobilising marker gene *asfA* and the sulfate ester mobilising *atsA* gene were analysed by generating respective clone libraries of the static and rotating treatments of hyphosphere and rhizosphere soil with *A. stolonifera* as host plant.

The *asfA* gene was amplified with primers asfAF1all (Gahan & Schmalenberger, 2015) and asfBtoA (Schmalenberger & Kertesz, 2007) (supplementary Table S1). A touchdown PCR was carried out under the following conditions: initial denaturation at 98 °C for 3 min, 10 cycles of 98 °C denaturation (10 s), 65-55 °C touchdown (15 s), 68 °C extension (40 s), plus 25 further cycles at 58 °C annealing. Final extension was carried out at 68 °C for 3 min. PCR was undertaken in 25 µL reactions containing; 1 X Terra PCR Direct Buffer (2 mM MgCl_2_),0.2 mM dNTP mix, 0.4 µmol of each primer, and 0.7 U of Terra PCR Direct Polymerase (ClonTech Europe, Saint-Germain-en-Laye, France).

The *atsA* gene was amplified using the AtsA-F1 AtsA-R1 primer pair (Ikoyi *et al*., 2020) (supplementary Table S1). PCR conditions were as follows: initial denaturation at 95 °C for 3 min, 40 cycles of 95 °C denaturation (60 s), 48 °C touchdown (60 s), and 72 °C extension (60 s). Final extension was carried out at 72 °C for 10 min. The PCR was undertaken using a Kapa 2G Robust PCR kit (Kapa Biosystems, Woburn, MA) in 25 µL reactions containing; 1 X Buffer A, 1 X Enhancer, 5% DMSO (Sigma-Aldrich), 2 mM MgCl_2,_ 0.2 mM dNTP, 0.5 µM of each primer and 0.5 U of the Kapa Robust polymerase.

The *asfA* and *atsA* PCR products were purified, quantified, and ligated into the cloning vector pJET1.2/blunt (CloneJet, Thermo Scientific) and pGEM^®^-T vector kit (Promega, Madison, Wisconsin, USA), respectively. The ligation and transformation steps were undertaken as per manual instructions. The ligation reactions were transformed into *E. coli* DH5α. In order to ascertain taxonomic diversity of recombinant plasmids containing an insert of the correct size, Restriction Fragment Length Polymorphism (RFLP) analysis was carried out on PCR amplicons using the restriction enzymes *AluI* and *RsaI* (5 U per reaction; Thermo Scientific) for 4.5 h at 37 °C. The digested DNA was run on a 2% agarose gel at 100 V for 40 min. Clones with a similar restriction pattern were classified as a single genotype using Phoretix 1D (Nonlinear Dynamics, Newcastle upon Tyne, UK). Unique genotypes with more than one representative were re-amplified and the purified PCR product was used for sequence identification (GATC Biotech). The sequences obtained were subjected to gene comparison using BLAST (Altschul *et al*., 1990). Sequences of *asfA* and *atsA* were imported into arb (Version 5.2) (Ludwig *et al*., 2004), translated into proteins and integrated into an established *asfA* phylogenetic trees (Schmalenberger *et al*., 2010) and *atsA* phylogenetic trees (this study). Nucleic acids were deposited in the nucleic acid archive under the accession numbers MT309545-MT309559 & MT380723 (*atsA*) and MT309560-MT309575 & MT372478-MT372479 (*asfA*).

### 2.1.1. Next generation sequencing

For Illumnia MiSeq 16S library preparation, community DNA from *A. stolonifera* was amplified using the primer pair 16SF and 16SR to target the V3 and V4 region yielding a 460 bp amplicon (Klindworth et al., 2012) (supplementary Table S1). The PCR were undertaken in 25 µL reactions with 0.5 U of Kapa HiFi Taq, 1 X PCR buffer with 1.5 mM MgCl_2_, 0.2 mM dNTPs each (all Kapa Enzymes, Woburn, MA), and 0.4 µM of each primer. A touchdown PCR protocol was used with the following cycling conditions: initial denaturation at 98 °C for 5 min, 20 cycles of 98 °C denaturation (45 s), 68-58 °C touchdown (60 s), 72 °C extension (60 s), plus 20 further cycles with an annealing temperature at 58 °C. The PCR product was purified using the GenElute PCR purification kit (Sigma-Aldrich, St. Louis, MO). The indexing PCR was carried out to attach the dual indices (Appendix D6.2) and Illumnia sequencing adapters using the Nextera XT Index Kit (Illumnia, San Diego, CA) in accordance with the manufacturer’s instructions with a modified annealing temperature of 63 °C. The indexing PCR product was purified (as before) and quantified using the Qubit dsDNA HS Assay Kit (Life Technologies, Carlsbad, CA) on a Qubit 2.0 Fluorometer (Life Technologies, Carlsbad, CA). The DNA concentrations of each sample were adjusted to 4 nM in 10 mM Tris pH 8.5 and 5 µL was used to mix aliquots for pooling libraries with unique indices. Amplicons were directly sequenced on an Illumnia MiSeq NGS platform (Illumnia, San Diego, CA) in line with protocols at the Teagasc, Moorepark sequencing centre.ee.

#### 2.1.2. Data analysis

Univariate analysis of EA-IRMS data, percentage root colonisation, and MPN data was carried out using IBM SPSS statistics 20 (Version 22.0; IBM, USA). DGGE fingerprinting gels were digitalised and band patterns analysed with the software package Phoretix 1D (Nonlinear Dynamics). Cluster analysis using UPGMA was carried out and obtained band pattern matrixes were exported for DCA and permutation tests (Monte-Carlo with 9,999 replicates) as described previously (Schmalenberger *et al*., 2010).

The following steps were undertaken to analyse the Illumnia NGS data; raw Illumina paired-end sequence reads were merged using Flash (Magoč & Salzberg, 2011) and quality checked using the split libraries script in Qiime (Caporaso *et al*., 2010). Quality checking is two phase; for joining reads a min overlap of 10 bp was expected; then for joined reads anything less than 150 bp and quality of 19 was removed. Reads were clustered into OTUs with a similarity cut off at 97% and chimeras were removed with the 64-bit version of USEARCH (Edgar, 2010). Subsequently, OTUs were aligned and a phylogenetic tree was generated within Qiime (Caporaso *et al*., 2010). Alpha and beta diversity analysis was also implemented within Qiime (Caporaso *et al*., 2010). Alpha diversity was determined using the Shannon and Simpson diversity indices and the *Chao*1 richness estimator. Beta diversity was analysed using PCoA undertaken with EMPeror (Vázquez-Baeza *et al*., 2013). Taxonomical assignments were reached using the 16S-specific SILVA database (Version 106). Sequences were deposited in the nucleic acid archive with the accession numbers (ERS4414312-ERS4414324).

## Results

### 3.1. Uptake of ^34^S from organo-^34^S enriched soil microcosms, ^34^S uptake from sulfatase treated & 1 µm mesh organo-^34^S microcosms

Organo-^34^S enriched soil was placed in AM selectively accessible mesh cores which were inserted into soil microcosms, this was undertaken to uncover the role of AM fungi in mobilisation and uptake of organo-S. Furthermore, experimental microcosms were created to uncover the role of AM in mobilisation of sulfonates and mesh cores were established that were inaccessible to AM hyphae to uncover the amount of ^34^S up-taken by the plant via potential leaching.

For the organo-^34^S enriched soil microcosms, all replicates of *A. stolonifera* at 3, 6, 9 (time of harvest) and 12 months (3 months post transfer) were subjected to EA-IRMS to determine organo-^34^S uptake and the percentage of ^34^S in the plant biomass. In addition, 3 *P. lanceolata* replicates per static and rotating treatment were selected for EA-IRMS at 3 and 6 months.

For *P. lanceolata*, total sulfur uptake was significantly increased (*P* < 0.05) for the static cores over the rotating cores at 3 months (0.23 vs. 0.20 %), but not at 6 months. Likewise, *A. stolonifera*, sulphur uptake increased significantly for the static cores for 3 and 6 months but not for month 9 (0.14, 0.23 and 0.23% for rotating cores and 0.16, 0.31 and 0.23 % for the rotating cores, respectively).

Very similar results were obtained for the ^34^S uptake. The δ^34^S_V-CDT_ (‰) values were significantly higher for the static cores for *Agrostis* and *Plantago* after 3 months of incubation when compared to the rotating core experiment. After 6 months this was only the case for *Agrostis* and after 9 and 12 months, no significant difference in δ^34^S_V-CDT_ was detected in the *Agrostis* plants (Figure 1 and S3). The δ^34^S_V-CDT_ (‰) values were relatively small at this time point (30 compared to 750 δ^34^S_V-CDT_ upon original analysis) which may explain the absence of a significant effect.

**Figure 1.**
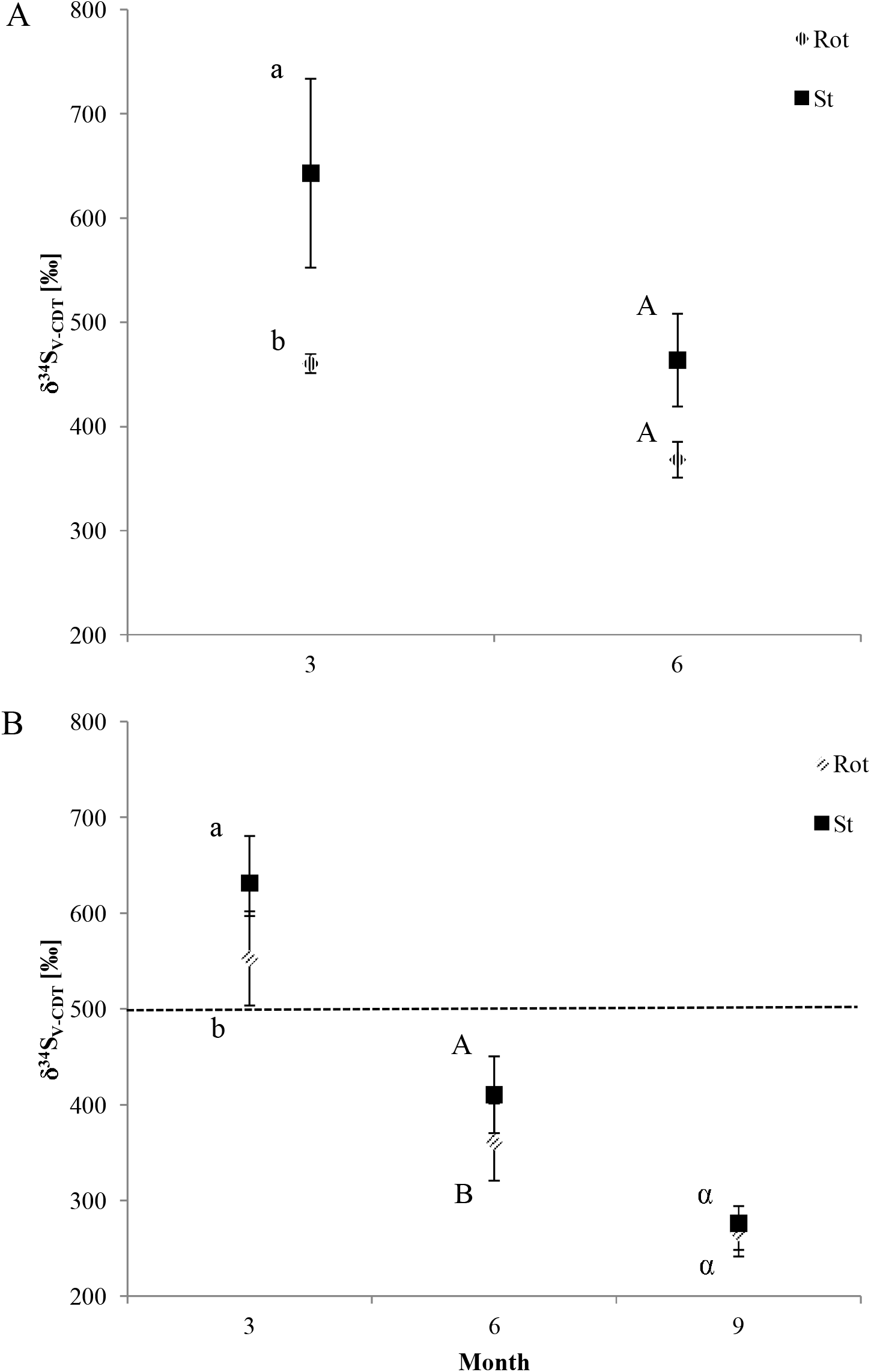
AM uptake of ^34^S from organo-^34^S following 3, 6 and 9 months of growth. A *= Plantago lanceolata*, B *= Agrostis stolonifera*, rotating (Rot) *=* severed hyphae, and static (St) = mycorrhizal. *P. lanceolata* was measured at 3 and 6 months, only (3 replicates per treatment). *A. stolonifera* was measured at 3, 6 and 9 months (all 6 replicates were analysed). Letters (A-B, a-b, α-β) indicate significant differences. The level of background (leaching) ^34^S uptake was measured at 3 months for *A. stolonifera* and is highlighted (dashed horizontal line).

In contrast, uptake of ^34^S for static and rotating core experiment after sulfatase treatments were not significantly different after 3 months (*P >* 0.05). The δ^34^S_V-CDT_ values of the rotating and static core experiment did not exceed the background leaching observed in the 1 µm mesh microcosms at 3 months (Figure 2).

**Figure 2.**
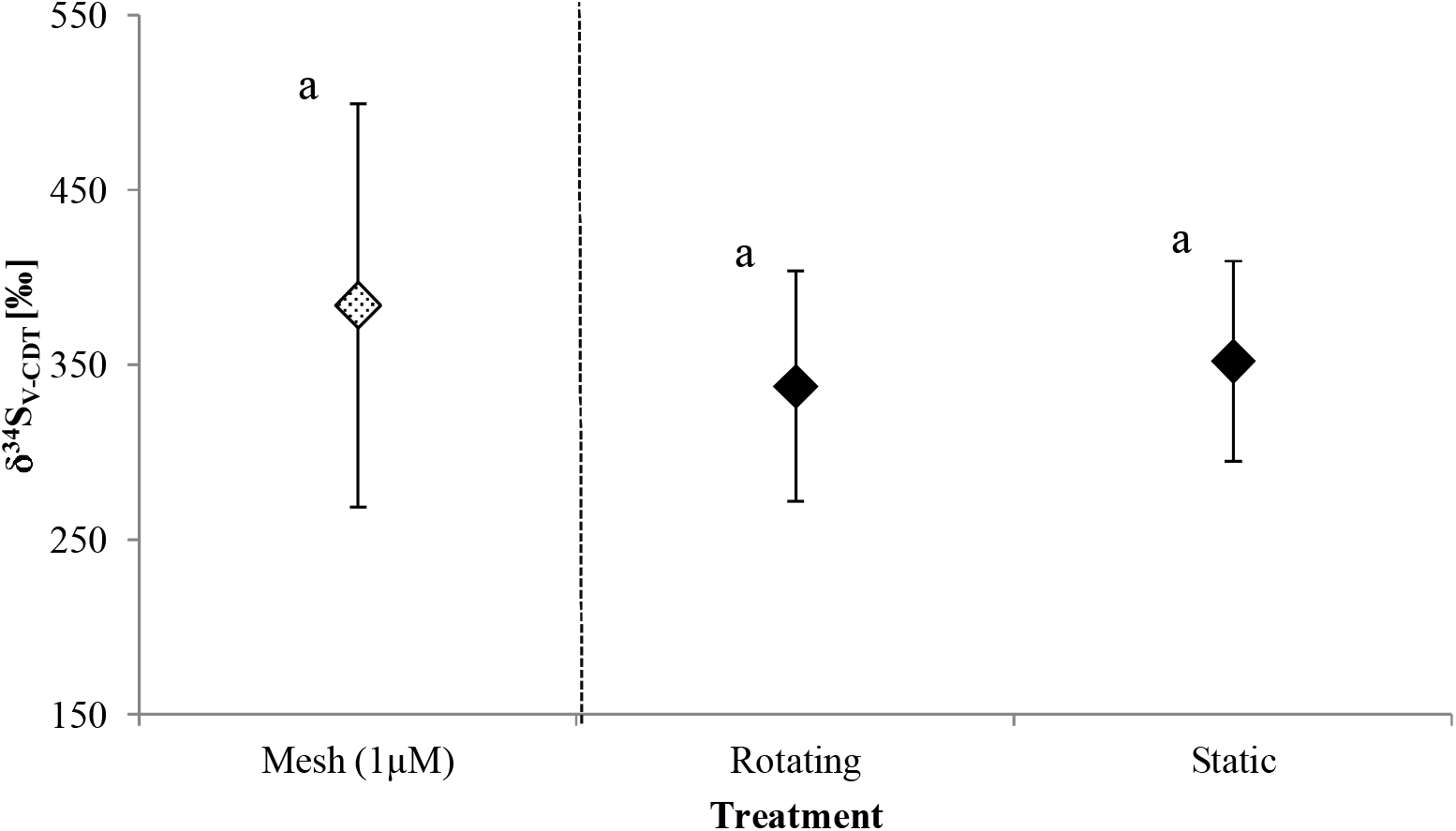
AM uptake of ^34^S from 1 µm mesh (not sulfatase treated, pattern fill) and sulfatase treated 35 µm mesh organo-^34^S microcosms (solid fill) with *Agrostis stolonifera* after 3 months of growth. Mesh (1 µm) was designed to quantify the effect of leaching. The sulfatase treated rotating *=* severed hyphae, and static = mycorrhizal treatments. Significant differences were not observed (a).

### 3.2. S K-edge X-ray absorption near edge spectroscopy

XANES analysis was undertaken to identify S oxidation states which include; reduced (sulfide, thiols), intermediate (sulfoxides and sulfonates) and oxidised (sulfates and sulfate esters) S species. XANES analysis revealed similar spectra for the static and rotating treatments, although, slightly more abundant reduced S forms were observed for the static treatments (supplementary Figure S4). The photon energy of reduced thiols, intermediate sulfonates and oxidised sulfate esters has been revealed to be 2474.4, 2480.2-2480.4, and 2481.6 eV, respectively (Schmalenberger *et al*., 2011). The photon energy for the corresponding S species in this study are larger due to a larger step size increment utilised in this study than that used to obtain the reference values (0.5 eV to 0.2 eV).

### 3.3. AM colonisation

The extent of root intracellular colonisation with characteristic AM structures was assessed for both *A. stolonifera* and *P. lanceolata* (Figure 3). Increased AC, VC and HC rates were observed for static over rotating treatment (*P* < 0.05). Rates of AC are increased over VC for both plants; this is as expected because the vesicle is not a characteristic structure of all AM species.

**Figure 3.**
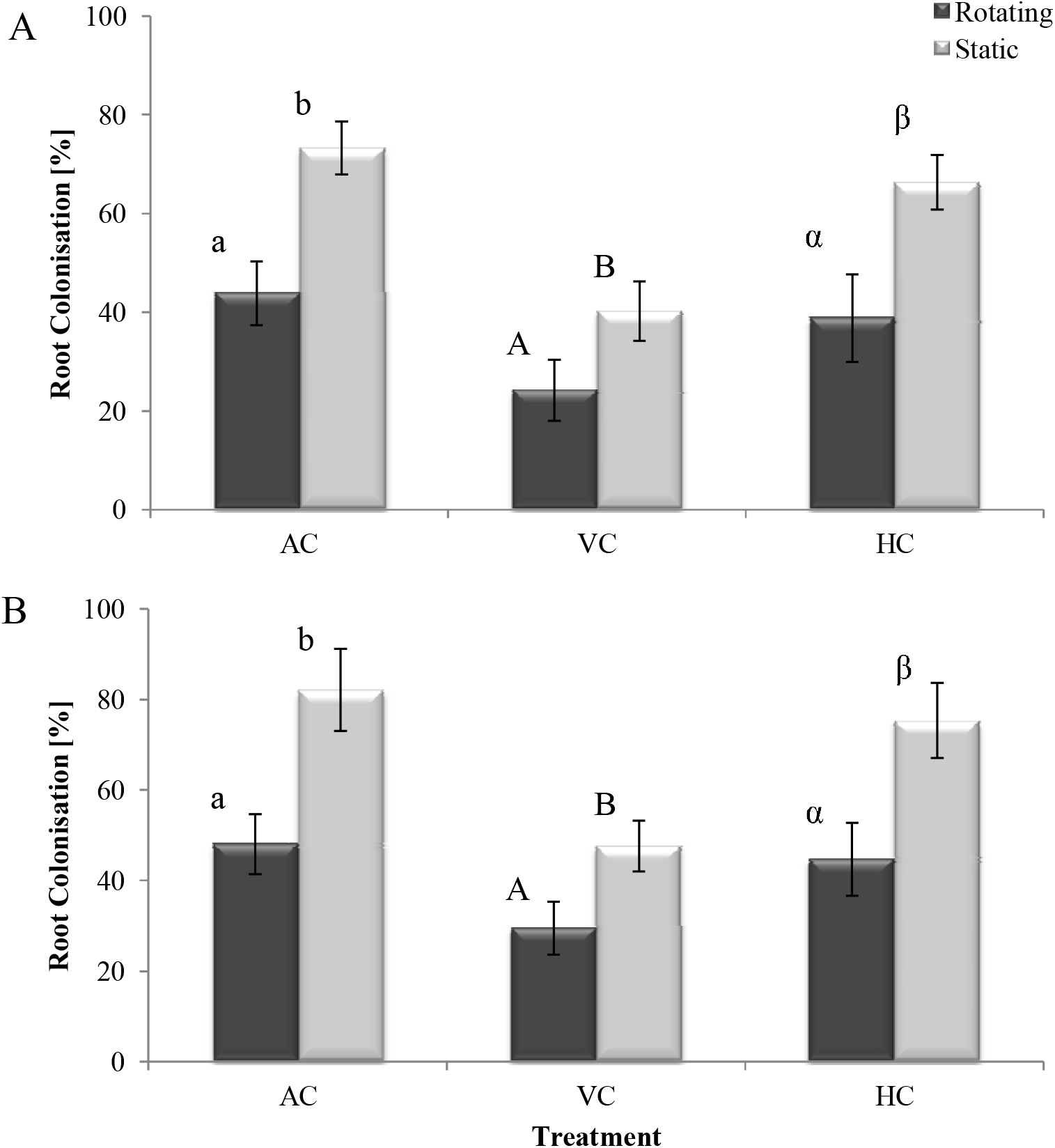
Arbuscular mycorrhizal root colonisation for *Agrostis stolonifera* (A) and *Plantago lanceolata* (B) in rotating and static microcosms. AC = Arbuscular Colonisation, VC = Vesicular Colonisation, and HC = Hyphal Colonisation. Letters (A-B, a-b, α-β) indicate significant differences.

### 3.4. Quantification of cultivable heterotrophic microbes

The total cultivable heterotrophic, sulfonate and polymeric sulfonate mobilising bacterial communities were quantified and compared to ascertain the impact of the presence of an intact AM symbiosis had on the magnitude of the respective bacterial community. This was achieved via MPN g^-1^ analysis in the corresponding media for the hyphosphere and rhizosphere, static (mycorrhizal) and rotating (severed hyphae) treatments.

For both plants, cultivable heterotrophic (R2A) and sulfonate mobilising (MM2TS) bacterial communities were more abundant in the microcosms with the static treatment for both the static cores as well as the rhizosphere (roots) and when compared to the respective rotating treatment (*P* < 0.05) (Figure 4A, B). Additionally, for *A. stolonifera* the polymeric sulfonate mobilising (MM2LS) bacterial community was more abundant in the static treatment (rhizopshere and cores) over the respective rotating treatment (*P* < 0.05) (Figure 4A). For *P. lanceolata*, the polymeric sulfonate mobilising bacterial abundance was not significantly different in the differently treated cores and rhizospheres with the exception of the rhizosphere for the rotating treatment, which was significantly lower in abundance (*P* < 0.05) (Figure 4B).

**Figure 4.**
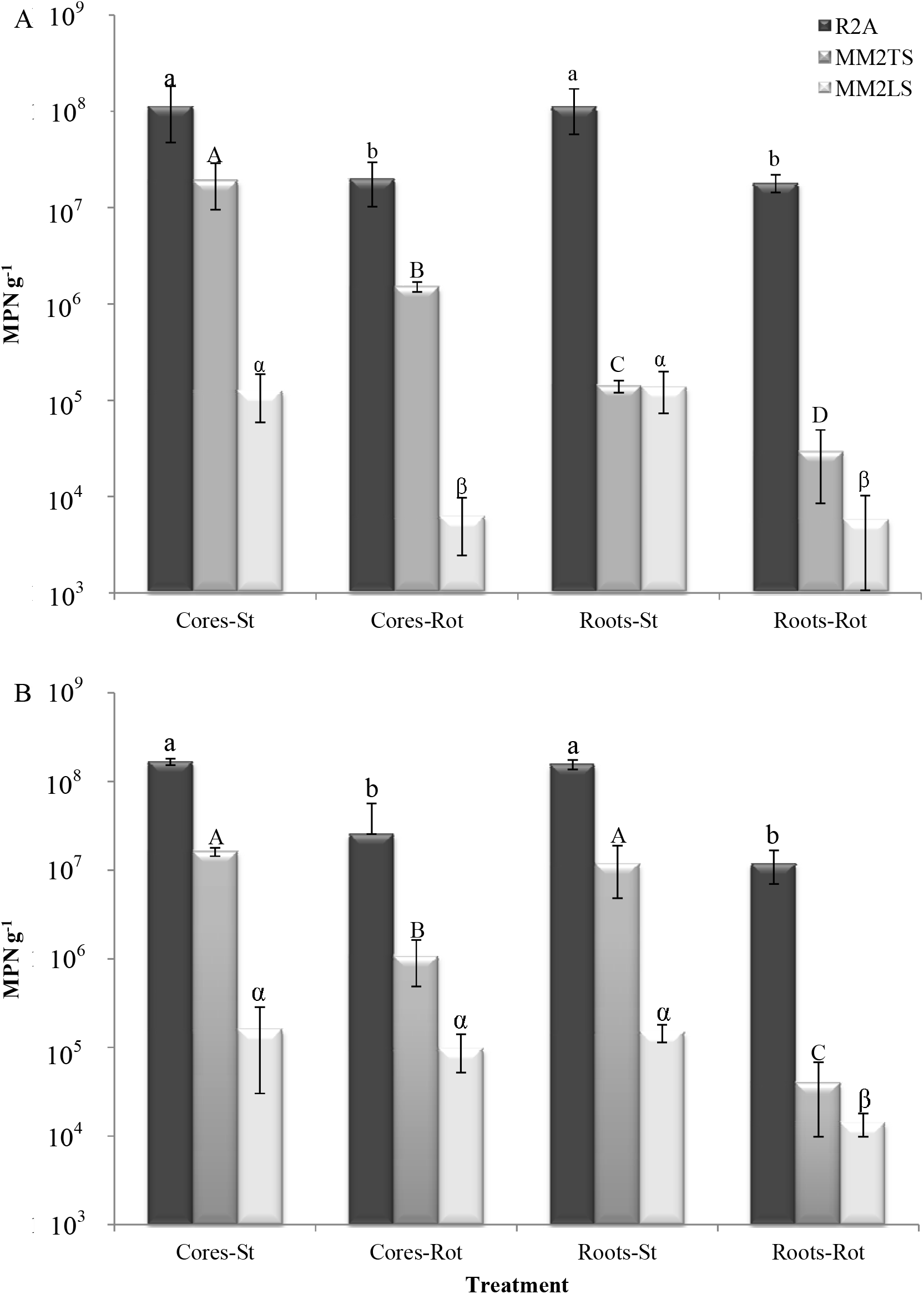
Most probable number (MPN) analysis for *Agrostis stolonifera* (A) and *Plantago lanceolata* (B). R2A, MM2TS and MM2LS were used to quantify the total cultivable bacterial community, sulfonate mobilisers polymeric sulfonate mobilisers, respectively. Cores = hyphosphere, roots = rhizosphere, static (St) = mycorrhizal, and rotating (Rot) = severed mycorrhiza. Letters (A-C, a-b, α-β) indicate significant differences.

The total cultivable heterotrophic bacterial abundance was the highest for both plants, followed by the sulfonate mobilisers and was lowest for the polymeric sulfonate mobilising bacteria (Figure 4A, B). The bacterial abundance was not different between cores and rhizospheres for both plants (*P* > 0.05) with the exception of the sulfonate mobilisers in *A. stolonifera* microcosms which was more abundant for cores over the rhizosphere (*P* < 0.05) (Figure 4A).

### Community fingerprinting

Based on DCA analysis and Monte Carlo permutation tests of *A. stolonifera* and *P. lanceolata* Bacterial (Figure 5A, B), AM fungal (Figure 6A, B) and saprotrophic fungal communities were significantly different for both plant hosts (Figure 7A, B) for all experimental treatments (*P* < 0.05), with the exception of the rhizospheres of (static over rotating) for *A. stolonifera* (Figure 7A) (*P* > 0.05).

**Figure 5.**
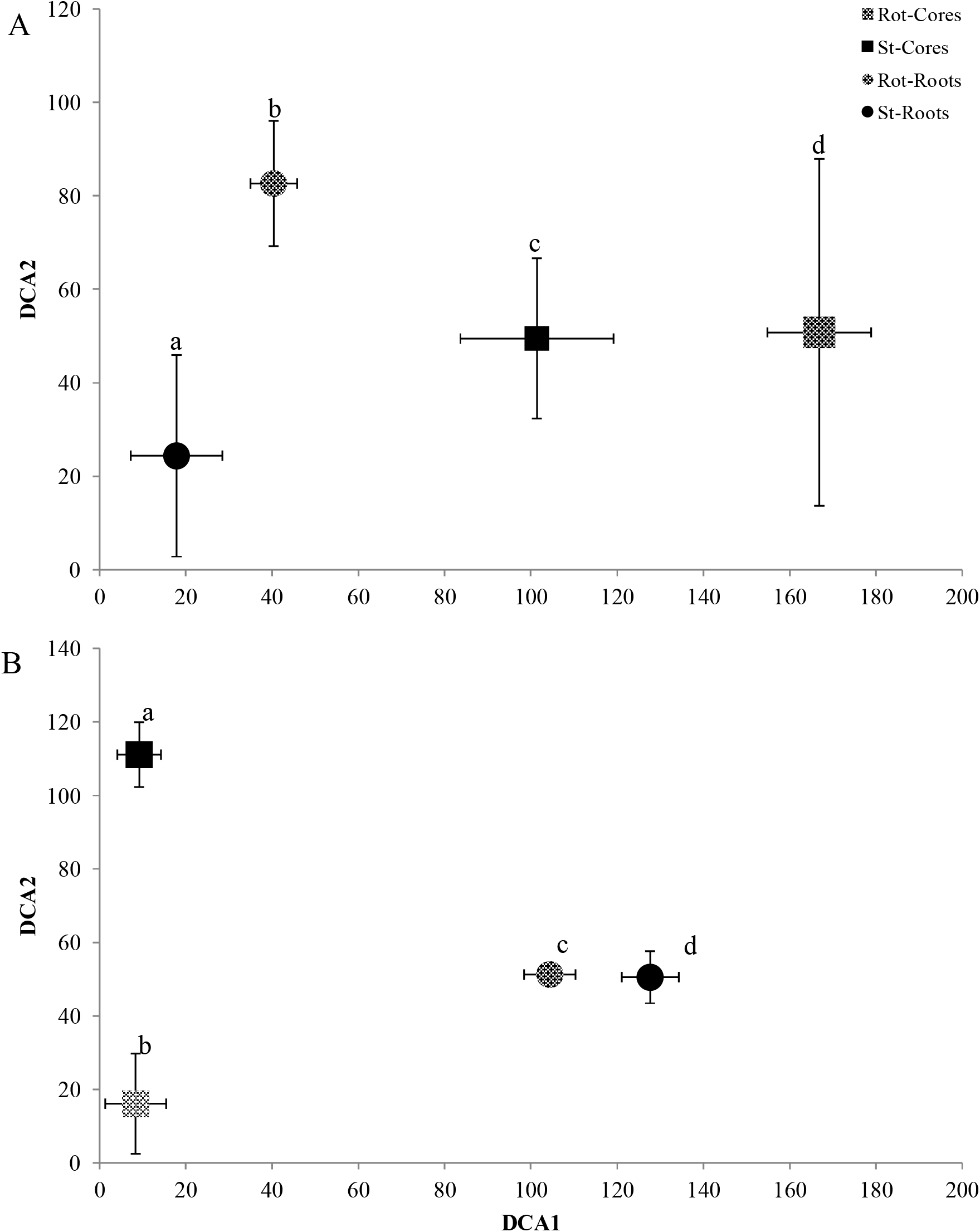
Detrended correspondence analysis (DCA) bi-plot of the 16S rRNA community fingerprint for *Agrostis stolonifera* (A) and *Plantago lanceolata* (B). CA1 = Axis 1, CA2 = Axis 2. Treatments; roots = circle, cores = square, static (st) = solid, rotating (rot) = pattern. Letters (a-d) indicate significant differences.

**Figure 6.**
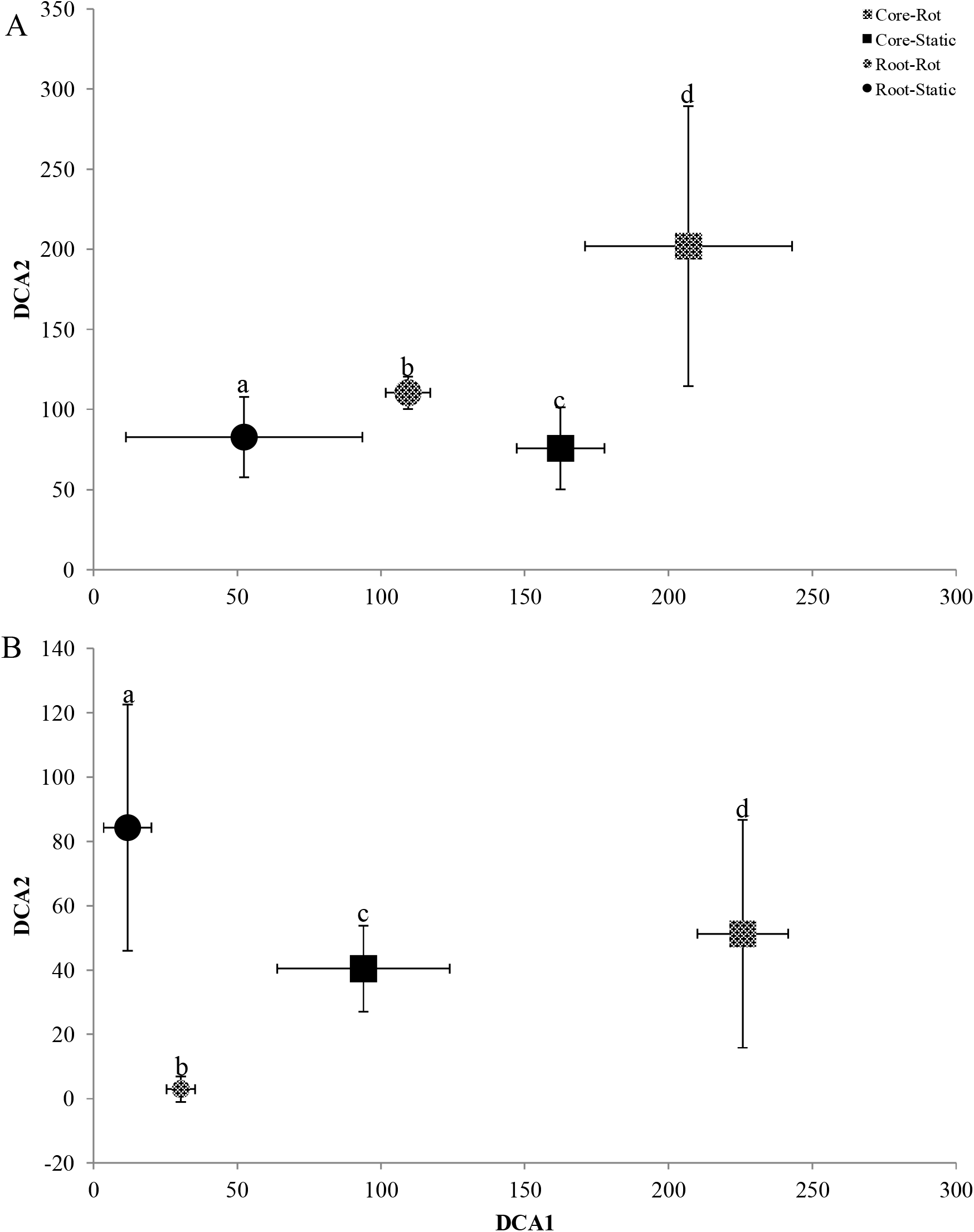
Detrended correspondence analysis (DCA) bi-plot of the 18S AM fungal community fingerprint for *Agrostis stolonifera* (A) and *Plantago lanceolata* (B). CA1 = Axis 1, CA2 = Axis 2. Treatments; roots = circle, cores = square, static (st) = solid, rotating (rot) = pattern. Letters (a-d) indicate significant differences.

**Figure 7.**
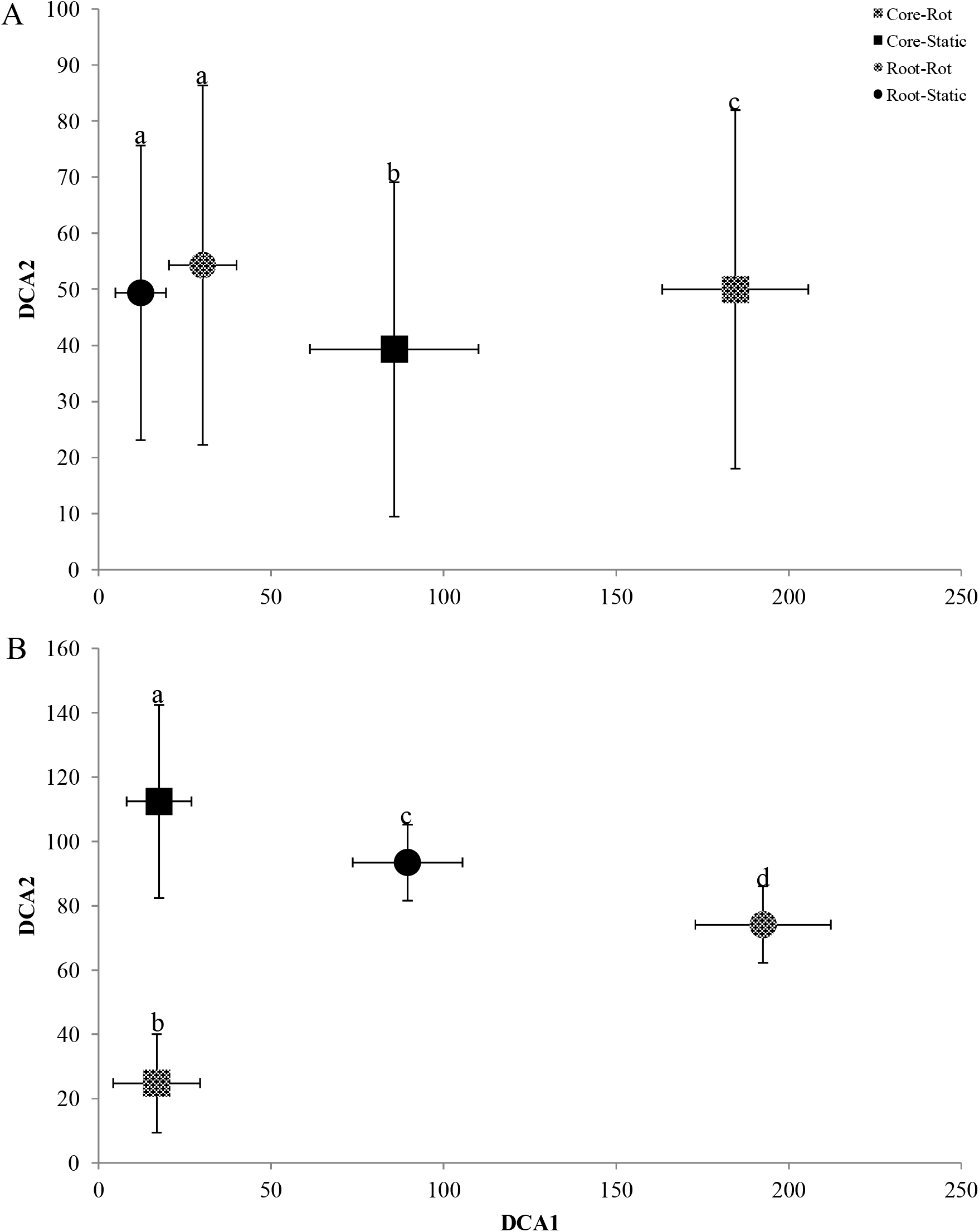
Detrended correspondence analysis (DCA) bi-plot of the fungal ITS community fingerprint for *Agrostis stolonifera* (A) and *Plantago lanceolata* (B). CA1 = Axis 1, CA2 = Axis 2. Treatments; roots = circle, cores = square, static (st) = solid, rotating (rot) = pattern. Letters (a-d) indicate significant differences.

#### 3.6. Arylsulfatase activity

Sulfate ester mobilising activity was measured in the hyphosphere and rhizosphere of the static (mycorrhizal) and rotating (severed hyphae) treatments. For both plants, arylsulfatase activity was higher for the rhizosphere than the cores and, additionally, for the rhizospheres from the static over the rotating treatment (*P* < 0.05). For *A. stolonifera*, 19.1 (± 1.09) and 11.8 µg (± 1.90) of *P*-nitrophenol was released in the rhizosphere from the static and rotating treatments, respectively. For the static and rotating cores this was lower at 5.5 (± 1.22) and 8 µg (± 2.23), respectively and overlapped. For *P. lanceolata*, 23.4 (± 2.25) and 17 µg (± 2.30) of *P*-nitrophenol was released from rhizospheres of the static and rotating treatments, respectively. This was significantly higher when compared to the static and rotating cores at 2.5 (± 0.50) and 2.5 µg (± 0.30) that overlapped as well (*P* > 0.05) (supplementary Figure S5).

### 3.7. Diversity of asfA and atsA gene

In order to ascertain the diversity of sulfonate and sulfate ester mobilising bacterial communities in the hyphosphere and rhizosphere of static (mycorrhizal) and rotating (severed hyphae) treatments, clone libraries of the aromatic sulfonate mobilising *asfA* and sulfate ester mobilising *atsA* gene were generated.

Screening of 200 *asfA* clones (50 each per treatment) revealed 43 OTUs in total; 12 in static cores, 10 in rotating cores, 12 in the rhizosphere of the static and 15 in rhizosphere of the rotating treatments. Only 2 OTUs occurred in all treatments. Both OTUs were the most abundant composing 23% and 12% of the sulfonate mobilising population analysed. Library coverage was calculated (Schmalenberger *et al*., 2007) to be between 92 and 96%. Of these 43 OTUs, 18 OTUs had more than three representatives and were subjected to DNA based sequence identification. Preliminary identification of the sequences was carried out with BLAST (Altschul *et al*., 1990). All sulfonate mobilising bacterial marker gene fragments were found to belong to the phylum Proteobacteria (Table 1).

**Table 1.** Taxonomic assignment of representative desulfonating operational taxonomicunits (OTUs) derived from clonal *asfA* DNA sequence analysis from hyphosphere cores (C) and rhizosphere roots (R) from static (S) = mycorrhizal and rotating (R) = severed mycorrhizal hyphae treatments.

| OTU ID | Affiliation | Abundance per treatment |  |  |  | Similarity [%]<br>DNA/Protein |
| --- | --- | --- | --- | --- | --- | --- |
|  |  | CS | CR | RS | RR |  |
| RR2 | α-Proteobacteria | | | | 8 | 70/84 |
| RR8 | α-Proteobacteria | | | | 4 | 71/68 |
| RS34 | α-Proteobacteria | | | 3 | | 70/68 |
| CS37 | Rhizobiales | 5 | 11 | 2 | 6 | 72/66 |
| CR11 | Rhizobiales |  | 6 |  |  | 72/66 |
| CS16 | Rhizobiales | 29 | 10 | 5 | 2 | 72/66 |
| RR6 | Rhizobiales |  |  |  | 7 | 71/74 |
| RS6 | Comamonadaceae |  |  | 13 |  | 72/76 |
| RR34 | Comamonadaceae |  |  |  | 3 | 71/71 |
| RS4 | Comamonadaceae |  |  | 4 |  | 78/92 |
| RS40 | Comamonadaceae |  |  | 6 |  | 82/94 |
| CR38 | *Stenotrophomonas* |  | 8 |  |  | 65/88 |
| CS12 | *Paraburkholderia* | 3 |  |  |  | 99/98 |
| RR29 | *Variovorax* |  |  |  | 5 | 81/95 |
| CR8 | *Variovorax* |  | 4 |  |  | 88/99 |
| CR13 | *Variovorax* |  | 4 |  |  | 83/92 |
| RR22 | *Variovorax* |  |  |  | 4 | 82/86 |
| RS20 | *Variovorax* |  |  | 8 |  | 82/86 |

For further phylogenetic analyses, sequences were imported into arb (version 5.2), translated into proteins and incorporated into a phylogenetic tree ((Schmalenberger *et al*., 2010) supplementary Figure S6, S7). The two OTUs present in all clone libraries (CS16 and CS37) represented 35 % of the clones (Table 1) and clustered in a clade that contains the genus *Cupriavidus* (supplementary Figure S7).

Five OTUs associated to Variovorax and one OTU associated at the family level of the Commamonadaceae (Table 1) clustered within the *Variovorax* clade of the phylogenetic tree (supplementary Figure S6). Together, they represented 16 % of the *asfA* clones. None of the *Variovorax* associated clones were found in the static mesh cores, while 8 of the 200 clones were found within the rotating cores. The remaining 23 clones were found in the plant rhizosphere.

The *Burkholderia* clade harboured one OTU (CS12), while the *Polaromonas* clades contained 2 OTUs (RS4, RS6). Via BLAST, the former OTU was associated with the genus *Paraburkholderia*. Only recently, the genus Burkholderia has been split into two genera, with most of the environmental strains moved into the newly established genus of *Paraburkholderia* (Dobritsa & Samadpour, 2016). The clade that was associated with *Cupriavidus* contained 40 % of the *asfA* clones that included CS37 and CS16. Notably, 29 of the 46 clones of CS16 were found on fungal hyphae in static mesh cores (presumptive mycorrhizal). One OTU was found to be closely associated with *Stenotrophomonas* (CR38; BLAST and phylogenetic tree) that was only found in the rotating mesh core.

Clone libraries of the sulfate ester mobilising *atsA* gene amplicons from the hyphosphere and rhizosphere of the static and rotating microcosms with *A. stolonifera* as host plant were screened for diversity via RFLP. Screening of 160 clones (40 each per treatment) revealed 38 OTUs in total; 9 for static cores (5 overlapping), 14 for rotating cores (6 overlapping), 15 for static roots (5 overlapping), and 13 for rotating roots (5 overlapping). Library coverage (Schmalenberger *et al*., 2007) was at or above 85%. Of these 38 OTUs, 15 OTUs with more than three representatives were sequenced, translated into proteins using Gene Runner (Version 4.0.9.68), identified using BLAST (Table 2), imported into arb (version 5.2), and incorporated into a phylogenetic tree (supplementary Figure S8).

**Table 2.** Taxonomic assignment of representative desulfonating Operational Taxonomic Units (OTUs) derived from clonal *atsA* amino acid sequence analysis from hyphosphere cores (C) and rhizosphere roots (R) from static (S) = mycorrhizal and rotating (R) = severed mycorrhizal hyphae.

| OTU | Affiliation | Abundance per |  |  |  | Overall | Similarity |
| --- | --- | --- | --- | --- | --- | --- | --- |
| ID |  | treatment |  |  |  | abundance | [%] |
|  |  |  |  |  |  | [%] |  |
|  |  | CS | CR | RS | RR |  |  |
| 1 | Bacteroidetes |  | 2 | 6 | 12 | 12.5 | 48 |
| 7 | *Planctomyces* | 1 | 3 |  | 7 | 7.5 | 50 |
| 10 | *Schlesneria* |  | 2 |  | 2 | 3 | 67 |
| 19 | *Schlesneria* | 1 |  | 5 | 6 | 8 | 52 |
| 20 | *Acidobacteria* |  |  |  | 6 | 4 | 54 |
| 36 | *Acidobacteria* |  |  | 3 |  | 2 | 53 |
| 2 | *Chthoniobacter* |  | 2 | 1 | 2 | 3 | 67 |
| 14 | *Chthoniobacter* |  | 4 |  |  | 3 | 65 |
| 3 | *Planctomyces* | 3 | 2 | 2 | 1 | 5.5 | 50 |
| 16 | *Mucilaginibacter* | 3 |  |  |  | 2 | 59 |
| 4 | Rhodospirillales (order) |  | 6 |  |  | 4 | 51 |
| 5 | Rhodospirillales (order) | 7 | 2 |  | 1 | 6 | 54 |
| 18 | *Verrucomicrobia* | 6 |  |  |  | 4 | 57 |
| 23 | *Verrucomicrobia* | 3 |  |  | 1 | 2.5 | 57 |
| 33 | *Acidobacteria* |  |  | 6 |  | 4 | 54 |
| 30 | Ascomycota |  |  | 6 |  | 4 | 57 |

The dominating sulfate ester mobilising bacteria in possession of the *astA* marker gene were found to be associated with Planctomyces (13%), Bacteroidetes (12.5%) *Schlesneria* (11%), Acidobacteria (6%), *Chthoniobacter* (6%), *Mucilaginibacter* (2%), Rhodospirillales (order) (10%) and Verrucumicrobia (6.5%) (Table 2). One OUT, representing 4% of *atsA* sequences was associated with the fungal phylum Ascomycota. Differences in presence and abundance were observed across the four treatments. *Planctomyces* and *Schlesneria* were more abundant in the rhizosphere, particularly in the rotating treatments. *Chthoniobacter* and *Mucilaginibacter* were more abundant in the hyphosphere rotating treatments and static treatments, respectively. Rhodospirillales like species were predominantly isolated from the hyphosphere and were present in both static and rotating treatments. *Verrucomicrobia* were almost exclusively isolated from static roots and cores and were not detected in the rotating cores (Table 2).

### 3.8. Next generation sequencing

Using the *Chao* 1 index (Chao, 1984), the species diversity of the rotating and static cores (*A. stolonifera* as host plant) was not determined to be different at 4006 (± 1624) and 4774 (± 1242), respectively. The number of OTUs obtained from the rotating and static cores were determined to be 3955 (± 1644) and 4740 (± 1246), respectively. The rotating cores appeared to have a higher Simpson and Shannon index (0.97 and 7.4) than the static cores (0.87 and 6.6) (Simpson, 1949, Weaver, 1949).

PCoA (un-weighted, Unifrac distance matrix) and hierarchical clustering via un-weighted pair group method with arithmetic mean separated the rhizosphere and hyphosphere very clearly (Figure 8). The static and rotating root (rhizosphere) treatments clustered together; however, there was a partial separation between the static and rotating core (hyphosphere) treatments (Figure 8). This result corroborates the alpha diversity findings in that differences are apparent for the static mycorrhizal cores relative to the other treatments analysed.

**Figure 8.**
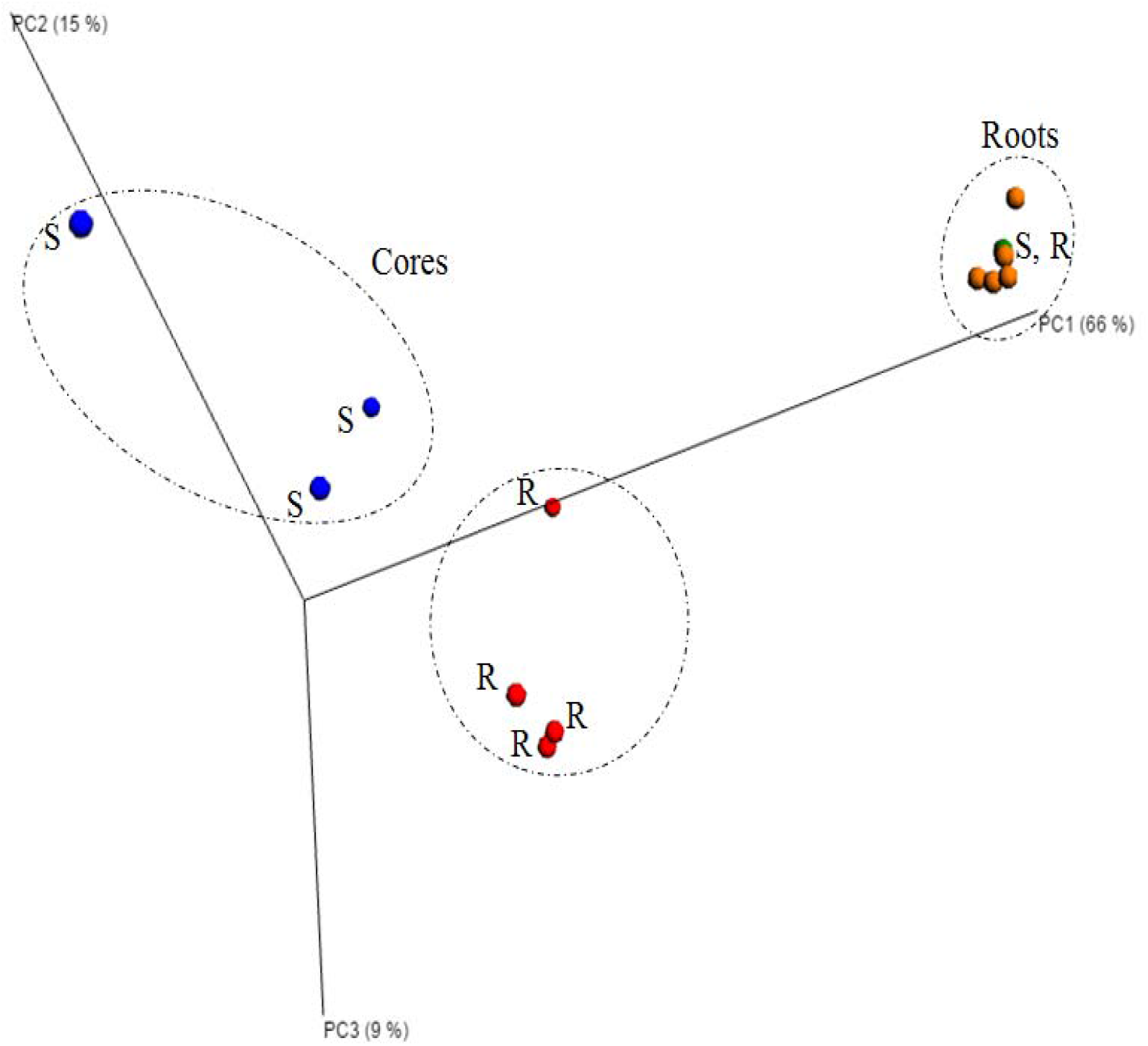
Principal component analysis (PCoA) of bacterial community sequences based on 16S rRNA amplicons from hyphosphere cores (C) (red (static) and blue (rotating)) and rhizosphere roots (R) (green (static) and orange (rotating)) of *Agrostis stolonifera*. Static (S) = mycorrhizal and rotating (R) = severed mycorrhizal hyphae treatments. PCoA was calculated using an un-weighted Unifrac distance matrix and visualised with EMPeror.

Taxonomic analysis of sequences retrieved from the hyphosphere and rhizosphere (static and rotating treatments) revealed that the Proteobacteria were the dominant phylum present for all treatments. The Proteobacteria were increased in abundance for both static cores and roots, at 28% and 31% compared to 22% and 27% for rotating cores and roots, respectively (supplementary Figure S9). Interestingly, all sulfonate mobilising *asfA* clones obtained belonged to the Proteobacteria.

Bacterial phyla that were dominant in the rhizosphere treatments included the Acidiobacteria (16-20%), Verrucomicrobia (9-10%), Chloroflexi (3-4%), Cyanobacteria (1-1.5%), Gemmatimonadetes (2.5-3%), and Nitrospirae (2%). Bacterial phyla that were dominant in the hyphosphere treatments included; Planctomycetes (34% and 21.4%) which compose 24% of the sulfate ester mobilising *atsA* gene community (see above), Firmicutes (2.5-3.5%), and Bacteroidetes (2-4%). The Planctomycetes, Proteobacteria and Verrucomicrobia were the only phyla that were more abundant in the static mycorrhizal treatments.

A number of phyla were more abundant for the rotating treatments which included; Firmicutes, Cyanobacteria, Chloriflexi and Acidiobacteria. Additionally, specifically for the mycorrhizal cores the Bacteroidetes and Actinobacteria were more abundant for the rotating treatments. These results corroborate the alpha diversity analysis by demonstrating a trend towards reduced diversity and evenness with evident dominance of certain bacterial phyla for the static mycorrhizal treatments.

At the family level, differences within the Proteobacteria were apparent between the core and root treatments (static, rotating) (Table 15). Families such as *Hyphomicrobiaceae* and *Bradyrhizobiaceae* were increased in abundance in the rhizosphere. While in the hyphosphere *Rhizobiaceae, Burkholderiaceae, Rhodocyclaceae, Comamonadaceae* and *Xanthomonadaceae* were the most prevalent.

For the core treatments, the dominating Proteobacteria for the static mycorrhizal cores were *Comamonadaceae* and *Xanthomonadaceae* composing 4.21% and 22.52% of the bacterial community, respectively. The remaining Proteobacteria families *Rhizobiaceae, Burkholderiaceae*, and *Rhodocyclaceae* were present in equal abundance for both static and rotating core treatments. Families that were prevalent in the hyphosphere (cores) and absent or occurred in low abundance in the rhizosphere (roots) included Planctomycetes *Planctomycetaceae*, Actinobacteria: *Micromonosporaceae*, Bacteroidetes: *Cytophagaceae* and *Porphyromonadaceae*, and Firimicutes: *Bacillaceae. Micromonosporaceae, Cytophagaceae* and *Porphyromonadaceae* were present/more abundant in the rotating over the static core treatments potentially as a result of the increased alpha diversity observed for the former (Table 3).

**Table 3.** Relative abundance (%) of major bacterial phyla and families in 16S rRNA gene fragment amplicon libraries generated using DNA from hyphosphere cores (C) and rhizosphere roots (R) of *Agrostis stolonifera* and static (S) = mycorrhizal (solid) and rotating (R) = severed mycorrhizal hyphae (pattern) treatments. nd = not detected.

| Phylum | Family | CR | CS | RR | RS |
| --- | --- | --- | --- | --- | --- |
| Proteobacteria | Betaproteobacterium (class) | nd | nd | 1.10 | 1.07 |
|  | uncultured bacterium | nd | nd | 1.02 | 1.75 |
|  | *Hyphomicrobiaceae* | nd | nd | 0.91 | 1.14 |
|  | uncultured bacterium | nd | nd | 1.79 | 1.62 |
|  | *Bradyrhizobiaceae* | nd | nd | 2.45 | 2.97 |
|  | *Rhizobiaceae* | 1.18 | nd | nd | nd |
|  | *Burkholderiaceae* | 1.25 | 1.29 | nd | nd |
|  | *Rhodocyclaceae* | 1.26 | 1.41 | nd | nd |
|  | *Comamonadaceae* | 1.57 | 4.21 | 1.21 | 1.03 |
|  | *Xanthomonadaceae* | 8.29 | 22.52 | nd | nd |
| Planctomycetes | uncultured soil bacterium | 1 | nd | nd | nd |
|  | uncultured bacterium | 18.48 | 18.73 | 1.44 | 1.19 |
|  | *Planctomycetaceae* | 4.03 | 2.97 | 2.00 | 1.98 |
|  | *Planctomycetaceae* | 1.94 | 1.23 | nd | nd |
|  | uncultured soil bacterium | 1.54 | 1.20 | nd | nd |
|  | Planctomycetales (order) | 1.12 | nd | nd | nd |
|  | Planctomycetales (order) | 1.12 | nd | nd | nd |
| Acidobacteria | uncultured bacterium | nd | nd | 1.39 | 1.45 |
|  | *Bryobacter* | nd | nd | 1.10 | 0.84 |
|  | uncultured bacterium | nd | nd | 2.32 | 1.47 |
|  | uncultured bacterium | 3.02 | 1.81 | 2.78 | 2.73 |
|  | uncultured bacterium | 6.47 | 3.79 | nd | nd |
| Actinobacteria | *Acidothermaceae* | nd | nd | 1.31 | 1.90 |
|  | *Micromonosporaceae* | 9.17 | 1.38 | nd | nd |
| Verrucomicrobia | Chthoniobacterales (order) | nd | nd | 6.01 | 6.02 |
|  | uncultured bacterium | nd | nd | 1.55 | 1.32 |
| Bacteroidetes | *Cytophagaceae* | 1 | nd | nd | nd |
|  | *Porphyromonadaceae* | 1.37 | nd | nd | nd |
| Nitrospirae | Nitrospirales (order) | nd | nd | 1.10 | 1.44 |
| Gemmatimonadetes | *Gemmatimonadaceae* | nd | 1.65 | 2.32 | 2.19 |
| Firmicutes | *Bacillaceae* | 1.89 | 1.78 | nd | nd |
| No blast hit |  | 15.14 | 17.80 | 18.87 | 19.26 |

Differences in familial community composition were observed for the Acidiobacteria with 2 uncultured bacteria and Bryobacter occurring only for the root treatments. Additionally, the Actinobacterium: *Acidothermaceae*, Verrucomicrobia: *Chthoniobacterales* and an uncultured bacterium, and the Nitrospirae: *Nitrospirales* occurred only for the root treatments. There was not a difference in familial community composition for the static and rotating root treatments which corroborates the results obtained with the alpha and beta diversity analysis (Table 3).

## 4. Discussion

AM fungi are highly beneficial for plant growth and have been shown to be involved in nutrient mobilisation using the organically bound labelled isotopes P and N (Hodge *et al*., 2001, Johnson *et al*., 2001, Hodge & Fitter, 2010). Additionally, AM fungi have been shown to promote growth (Rowe *et al*., 2007) and harbour more abundant sulfonate mobilising bacterial communities (Gahan & Schmalenberger, 2015). While little evidence exists to suggest AM fungi directly mobilise organically bound S, improved S uptake has been observed in their presence (Allen & Shachar-Hill, 2009) potentially as a result of bacterial metabolites improving ERH growth (Vilarino *et al*., 1997). This study aimed to illustrate that the presence of an intact AM symbiosis increases uptake of organo-S uptake via interactions with organo-S mobilising microbes.

Uptake of ^34^S from organo-^34^S enriched soil was increased in the static systems at 3 months for both *A. stolonifera* and *P. lanceolata* and 6 months for *A. stolonifera*. Plant S demand is dependent both on plant species and stage of development with increased demand observed during periods of vegetative growth and seed development (Leustek & Saito, 1999). When S demand is high plant SO_4_^2-^ transporters are up-regulated for rapid uptake in the rhizosphere leading to a SO_4_^2-^ depletion zone (Buchner et al. 2004). In this zone, bacterial desulfurisation of organo-S is induced via AM influenced rhizodeposition (Kertesz and Mirleau 2004). When S demand is lessened at later growth stages, it is no longer viable to expend C to stimulate proliferation of organo-S mobilisers and the AM role may be minimised. Additionally, it was observed that following treatment with an arylsulfatase enzyme to remove aromatic sulfate esters that AM fungi no longer increased ^34^S uptake from the organo-^34^S mesh cores. This fact may be a result of the relative availability of sulfate esters and sulfonates, many studies have demonstrated that in the short term S is almost exclusively mineralised from sulfonate pools (Ghani *et al*., 1992, Ghani *et al*., 1993, Zhao *et al*., 2006) which may suggest that the sulfonate-^34^S in this study was mineralised preceding mycorrhizal colonisation of the core. XANES analysis undertaken in this experiment uncovered that sulfate esters and sulfonates were both present in the soil and studies have shown that mobilisation of both organo-S species is important for optimisation of plant S supply (Freney *et al*., 1975, Freney, 1986). XANES analysis also revealed that the static mycorrhizal treatments were more abundant in reduced S species which indicates S uptake and incorporation into fungal biomass in the static cores. Indeed, more abundant reduced S has been associated with fungal biomass in the past (Schmalenberger & Noll, 2014).

The presence of an intact symbiosis in the static microcosms significantly increased root colonisation for both plants analysed. Tresender (2013) demonstrated that increased AM root colonisation improved both plant yield and nutrient content. The mechanism for which may include increased efficiency of nutrient mobilisation, uptake and transfer via characteristic mycorrhizal structures (Van Der Heijden *et al*., 1988, Klironomos & Hart, 2002). Additionally, an intact symbiosis alongside increased percentage root colonisation was shown to stimulate bacterial proliferation with more abundant cultivable heterotrophs, sulfonate and polymeric sulfonate mobilisers. This phenomena has been shown previously for cultivable heterotrophs and sulfonate mobilisers (Gahan & Schmalenberger, 2015). However, AM fungi are newly associated with stimulating bacterial populations capable of utilising high molecular weight polymeric sulfonates which may constitute an as yet un-quantified fraction of soil S (Kertesz *et al*., 2007). This ability involves either bacterial lignolytic or as yet uncharacterised extracellular sulfonatase activity (Kertesz *et al*., 2007).

Community fingerprinting was undertaken for the organo-^34^S microcosms and a bacterial, saprotrophic fungal and AM fungal community shift was observed for the static (mycorrhizal) and rotating (severed hyphae) treatments. This was as expected, as AM fungal symbiosis has been shown to alter microbial community composition in the hyphosphere and rhizosphere (Johansson *et al*., 2004, Gahan & Schmalenberger, 2015). A NGS approach was employed to identify diversity and taxonomy of bacterial species and this revealed reduced diversity and evenness for the static mycorrhizal cores in comparison to both the rhizosphere treatments and the rotating cores. Studies on bacterial diversity and abundance associated with AM hyphae have presented equivocal results in the past and the results obtained in this study were deemed to be a result of AM induced physico-chemical modification of rhizodeposition stimulating dominance of certain bacterial phyla and families (Andrade *et al*., 1997, Andrade *et al*., 1998, Scheublin *et al*., 2010). The Proteobacteria and Planctomycetes were both shown to be strongly stimulated to dominance in the static mycorrhizal treatments and the Proteobacteria are the dominant phylum of S mobilisers (Schmalenberger & Kertesz, 2007). Within the Proteobacteria, the *Xanthomonadaceae* were strongly stimulated by the presence of AM hyphae and this family includes *Stenotrophomonas* which was recently isolated directly from the hyphosphere and putatively has the ability to attach to and co-migrate with AM hyphae (Gahan & Schmalenberger, 2015). The *Comamonadaceae* were also stimulated specifically in the mycorrhizal treatments and this family includes; *Acidovorax, Hydrogenophaga, Polaromonas* and *Variovorax*, all of which have been associated with sulfonate mobilisation and the latter two specifically with AM hyphae (Schmalenberger & Kertesz, 2007, Fox *et al*., 2014, Gahan & Schmalenberger, 2015). This AM specific stimulation of sulfonate mobilising bacterial families may have contributed to the improved uptake of organo-S for the static mycorrhizal treatments.

Sulfate ester (arylsulfatase enzyme assay, *atsA* gene) and sulfonate mobilising activity (*asfA* gene) was significantly increased/altered by an intact AM symbiotic partnership. Arylsulfatase activity was higher in the rhizosphere of the mycorrhizal treatments and as uptake of ^34^S was increased only when sulfate esters constituted a fraction of the organo-^34^S pool, this activity presents a potential mechanism for this result. Furthermore, increased arylsulfatase activity may have been have been observed for the cores at an earlier stage of growth when S demand was higher. While AM fungi have not previously been shown to directly increase S uptake, previous studies have shown improved uptake in their presence and this may be a result of the C rich hyphosphere selecting for organo-S mobilising bacteria with the potential to improve nutrient supply (Banerjee *et al*., 1999, Knauff *et al*., 2003, Summerbell, 2005, Allen & Shachar-Hill, 2009).

Analysis of the diversity of the sulfate ester mobilising *atsA* gene revealed differences in presence and relative abundance of genera. The dominating sulfate ester mobilising genera in possession of the *atsA* marker gene were found to be associated to *Planctomyces* (20%), *Schlesneria* (17%), *Chthoniobacter* (6%), *Mucilaginibacter* (7.5%), Rhodospirillales (order) (10%), *Verrucumicrobia* (10.5%) and *Burkholderia* (4%). The *atsA* gene was most prevalent in the Planctomycetes which was also the most abundant phylum identified in the mycorrhizal cores via NGS which suggests that this functional population may be stimulated by AM fungal presence. However, as both *Planctomyces* and Verrucumicrobia (phylum) have only been identified from culture independent studies, their full rhizospheric functionality remains limited (da Rocha *et al*., 2009). The presence of *atsA* in *Burkholderia* is also very interesting as this genus has recently been identified as a sulfonate mobiliser with an intact *asfA* gene (Chapter 3 – Section 3.3.5) which suggests it may have dual functionality and contribute to the observed increased uptake of organo-^34^S.

In a pattern analogous to *atsA* gene diversity analysis, differences in presence and relative abundance of genera were also observed for the sulfonate mobilising community. *Cupriavidus* was the over-riding dominant desulfonating genus composing 58% of the sulfonate mobilising community in the static cores, this genus has been linked with improved P and N mobilisation in the past (Van Houdt *et al*., 2009, Yu *et al*., 2011) and may play an important role in overall plant nutrition. Additionally, *Burkholderia* and *Polaromonas* were dominant AM associated sulfonate mobilisers both of which have been isolated directly from AM hyphae and linked with sulfonate mobilisation in the past (Chapter 3 – Section 3.3.5). *Variovorax* species were also present but dominated the non-mycorrhizal treatments. In Chapter 3 of this thesis, *Variovorax* species were shown to dominate the control microcosms that did not receive AM inoculant while *Burkholderia* and *Polaromonas* were dominating species identified post inoculation (Section 3.3.5). The organo-^34^S microcosms in this study received inoculation with *R. irregularis* and this explains the similarity in species of sulfonate mobilisers obtained. Additionally, it suggests that while *Variovorax* species are important sulfonate mobilisers (Kertesz *et al*., 2007), post-inoculation other species increase in dominance. *Variovorax* species were also the dominant sulfonate mobiliser alongside correspondingly little mycorrhizal activity in an experiment conducted in Broadbalk, UK (Schmalenberger *et al*., 2008).

The results obtained demonstrate that an intact AM symbiosis increases uptake of ^34^S from organo-^34^S enriched soil at early stages of growth when S requirement is high. This may be a result of AM induced microbial community shifts leading to improved sulfate ester and sulfonate mobilising activity. In conclusion, AM symbiosis has potential to improve plant S uptake and reduce the requirement for unsustainable inorganic S fertilisation practices.

## 5. Acknowledgements

The authors would like to thank the staff of Teagasc, Johnstown Castle, for providing the soil used in this experiment, Jonathan Leake (University of Sheffield) to provide support for constructing the mesh cores, Christian Vogel and Wolfgang Pritzkow (BESSY II) for conducting the XANES analysis, and FP7 People (CIG no. 293429) for funding this project.

