## Supplementary figures, information and table for "The role of arbuscular mycorrhiza and organosulfur mobilizing bacteria in plant sulphur supply"

### Supplementary Material

#### Supplementary figures S1-S9

#### Supplementary information S1-S5

#### Supplementary table S1

Supplementary figures:

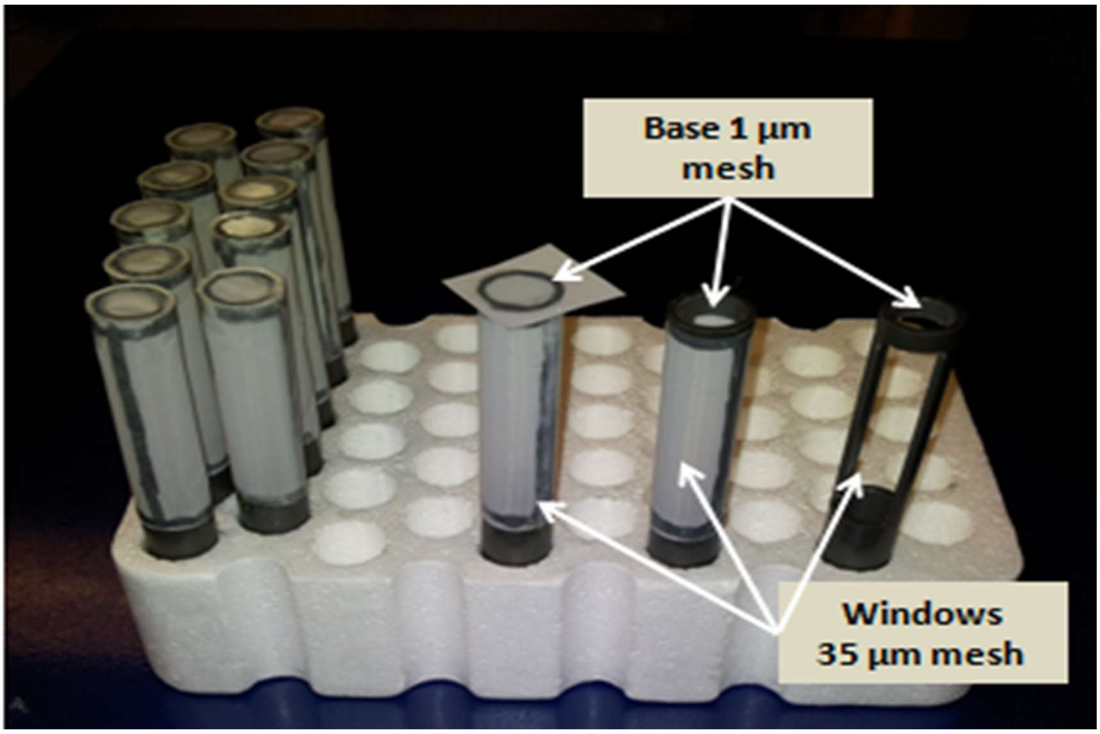


**Figure S1 – Construction of cores used to contain the $^{34}$S stable isotope enriched soil.**

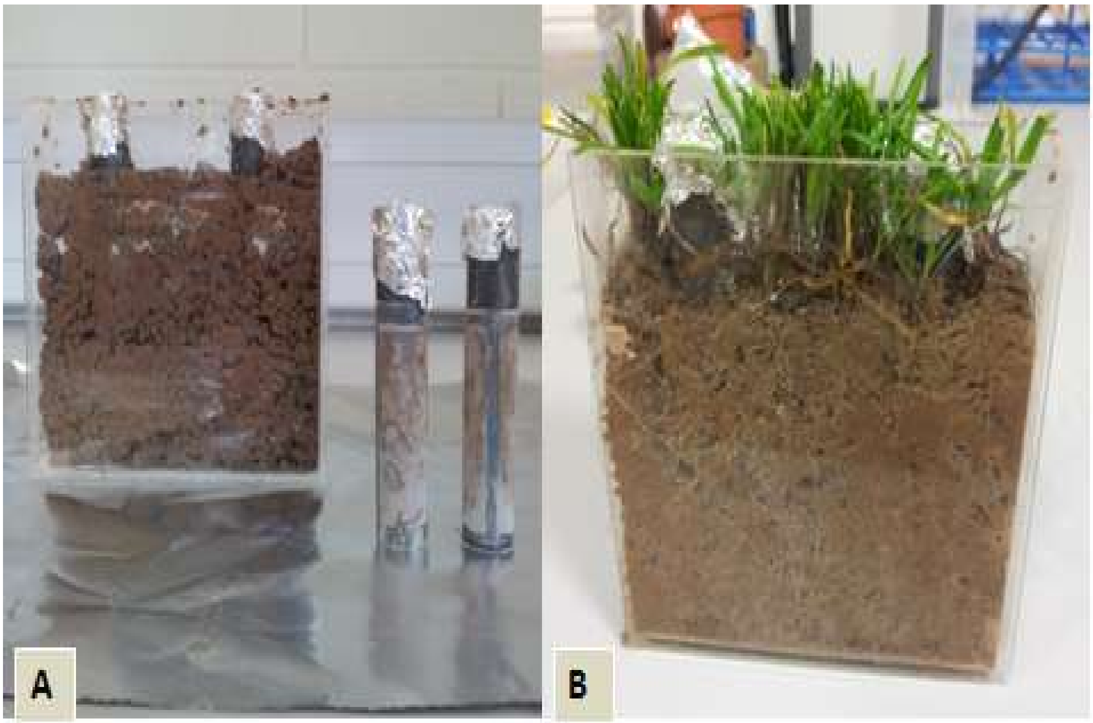


**Figure S2 – The soil microcosm systems with the organo-$^{34}$S enriched cores (A) and the actively growing system (B).**

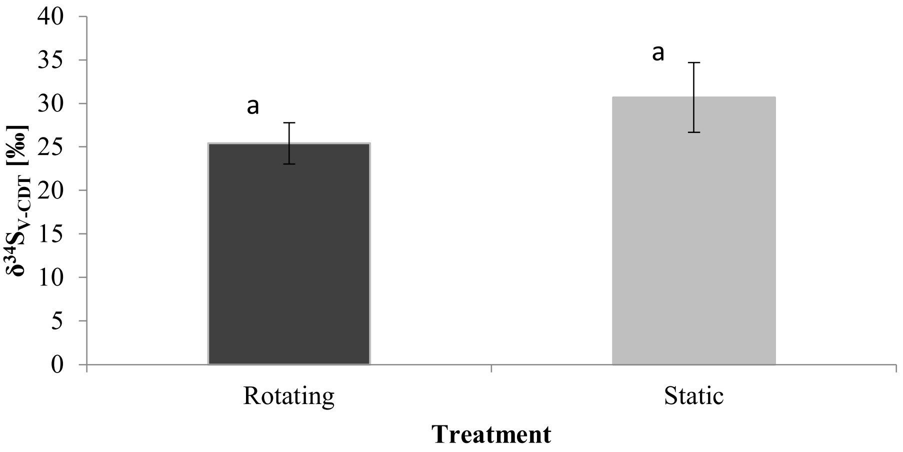


**Figure S1 – AM uptake of $^{34}S$ from organo-$^{34}S$ after 12 months (3 months) post transfer into new *Agrostis stolonifera* microcosms. Rotating = severed hyphae, Static = mycorrhizal. No significant differences (a).**

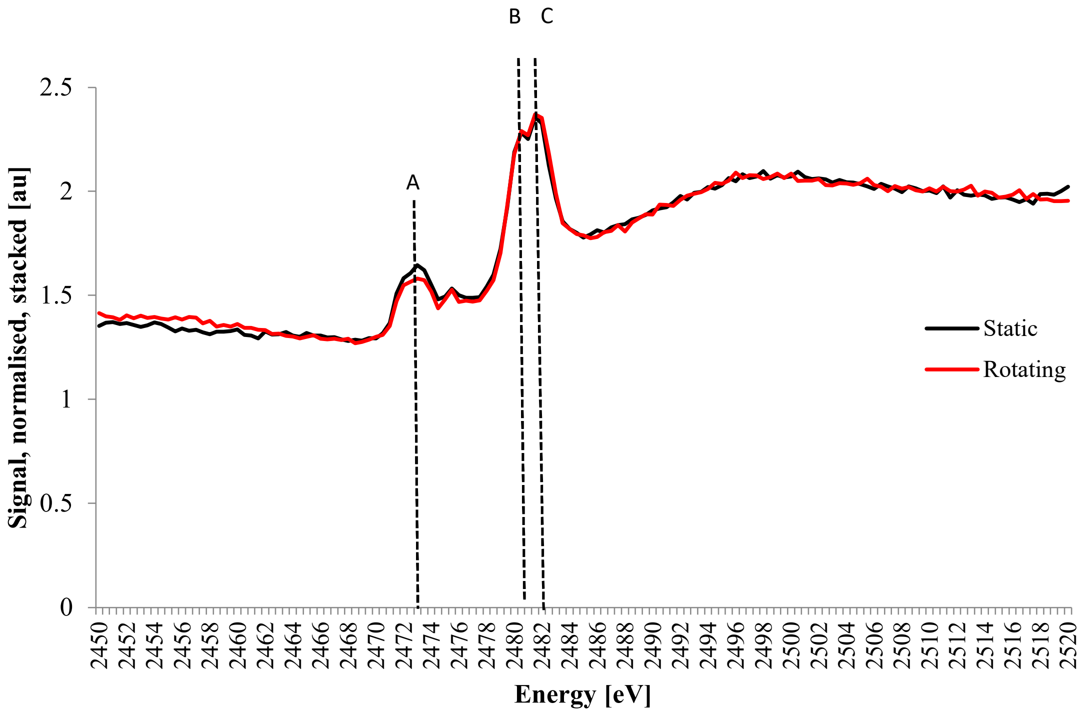


**Figure S4 – Normalised and stacked S K-edge spectra from static (black) and rotating (red) treatments with *Agrostis stolonifera* as host plant. Peak A represents reduced thiols, peak B represents intermediate sulfonates and peak C represents oxidised sulfate esters (Schmalenberger *et al.*, 2011, Schmalenberger & Noll, 2014).**

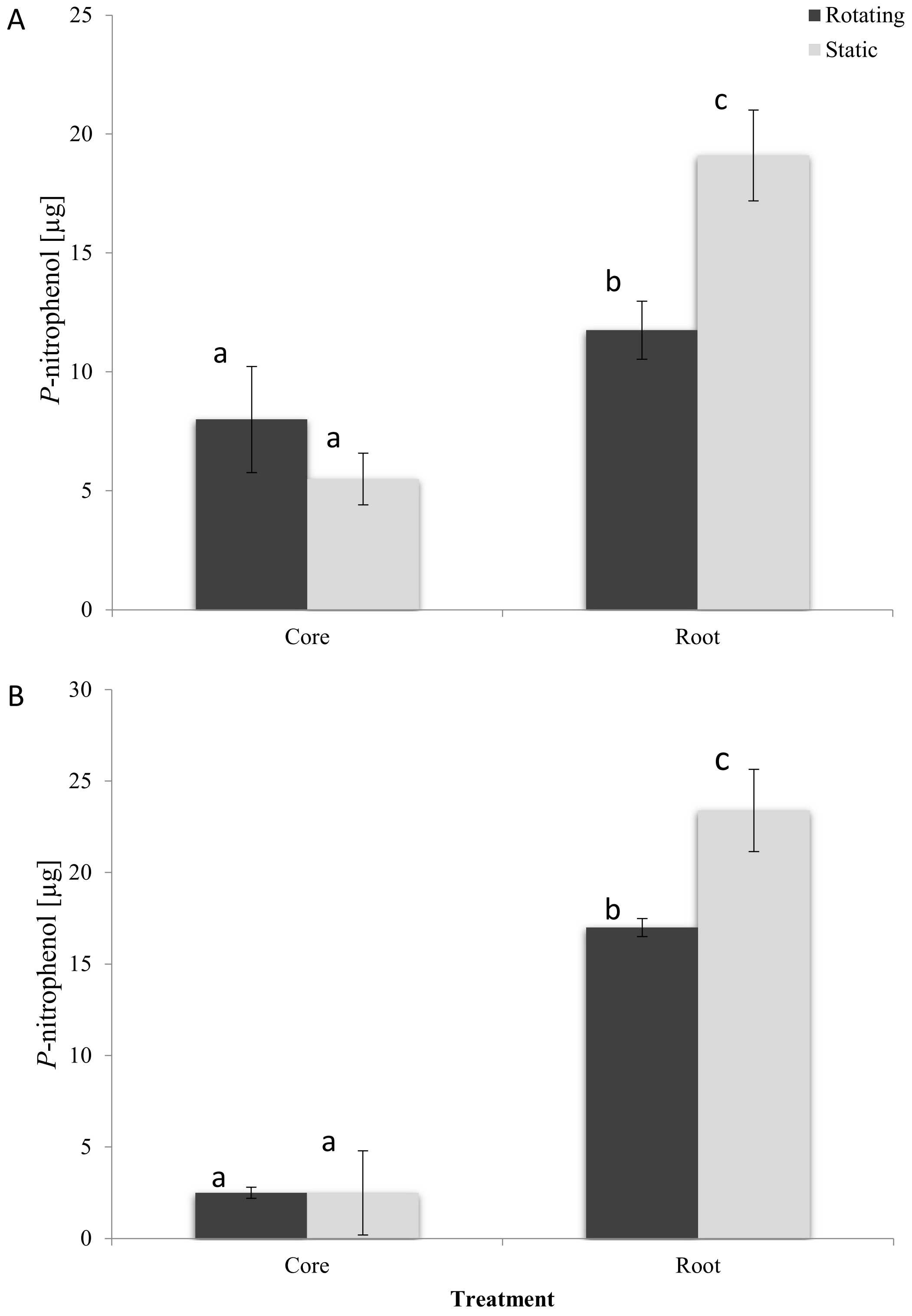


Figure S5 – Arylsulfatase activity for *Agrostis stolonifera* (A) and *Plantago lanceolata* (B) in organo-$^{34}S$ enriched soil microcosm systems. Rotating = severed hyphae, static = mycorrhizal, roots = rhizosphere, and cores = hyphosphere. Letters (a-c) represent significant differences.

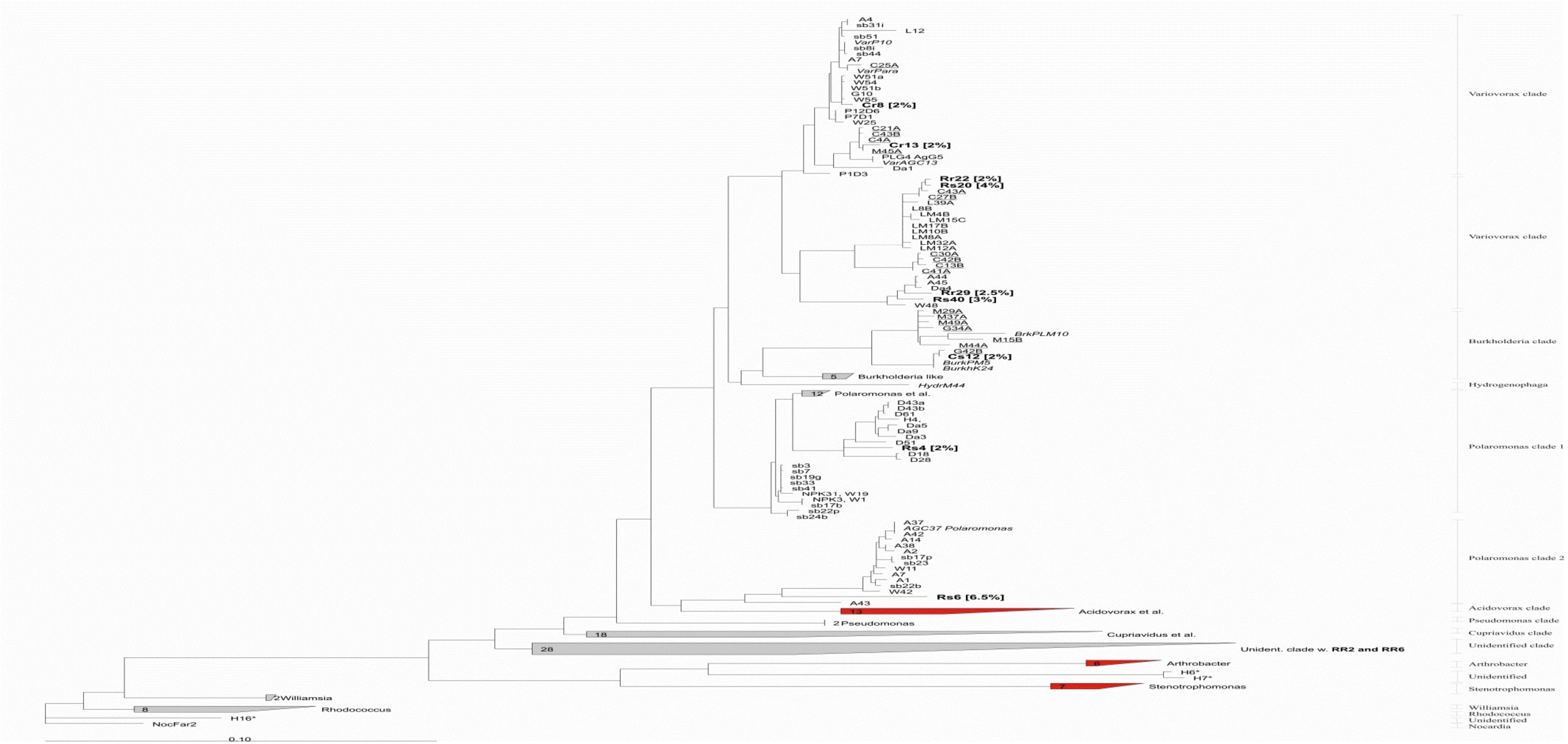


Figure S6 – Randomised axelerated maximum likelihood tree of truncated AsfA sequences of representative desulfonating operational taxonomic units (OTUs) derived from clonal *asfA* DNA sequence analysis from hyphosphere cores (C) and rhizosphere roots (R) from static (S) = mycorrhizal and rotating (R) = severed mycorrhizal hyphae treatments. Cultivated (*italics*) and molecular isolates from this study (bold) are highlighted. Molecular isolates from spring barley rhizospheres (sb; (Schmalenberger & Kertesz, 2007)), *Agrostis* grassland rhizospheres (CA; (Schmalenberger & Noll, 2010)), wheat rhizospheres from Broadbalk (W; (Schmalenberger *et al.*, 2008)), rhizospheres and soils from the Damma glacier forefield (D; DA; (Schmalenberger & Noll, 2010) ) and hyphosphere (H; (Gahan & Schmalenberger, 2015)) were isolated previously. Clades expanded include *Variovorax, Burkholderia,* and *Polaromonas*. Clades to be expanded in the figure below (Figure S7) are highlighted in red.

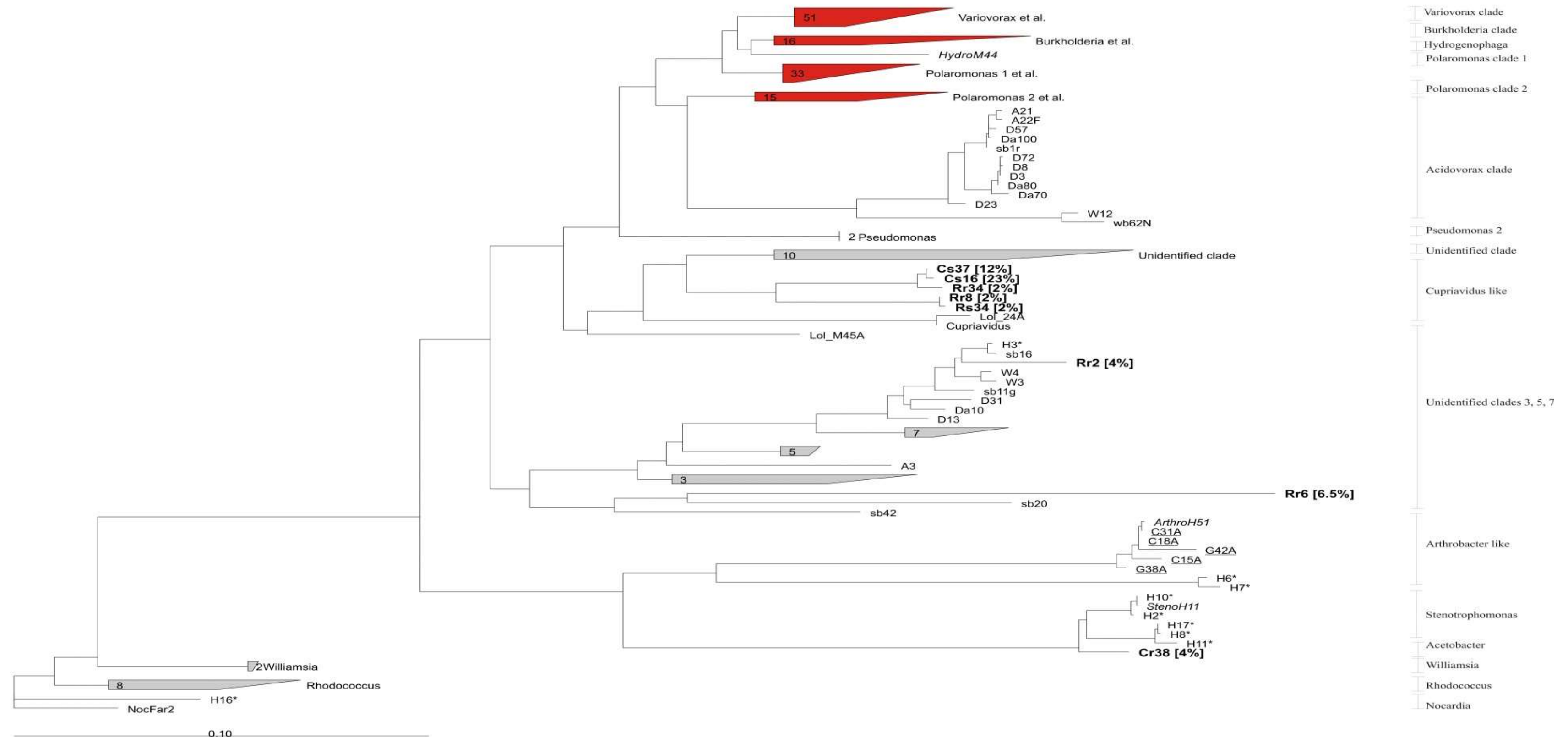


Figure 2 – Randomised axelerated maximum likelihood tree of truncated AsfA sequences of representative desulfonating operational taxonomic units (OTUs) derived from clonal *asfA* DNA sequence analysis from hyphosphere cores (C) and rhizosphere roots (R) from static (S) = mycorrhizal and rotating (R) = severed hyphae treatments. Cultivated (*italics*) and molecular isolates from this study (bold) are highlighted. Molecular isolates from spring barley rhizospheres (sb; (Schmalenberger & Kertesz, 2007)), Agrostis grassland rhizospheres (CA; (Schmalenberger & Noll, 2010)), wheat rhizospheres from Broadbalk (W; (Schmalenberger *et al.*, 2008)), rhizospheres and soils from the Damma glacier forefield (D; DA; (Schmalenberger & Noll, 2010)) and hyphosphere (H; (Gahan & Schmalenberger, 2015)) were isolated previously. Clades expanded include *Acidovorax, Arthrobacter* and *Stenotrophomonas*. Clades expanded in the preceding figure (Figure S6) are highlighted in red.

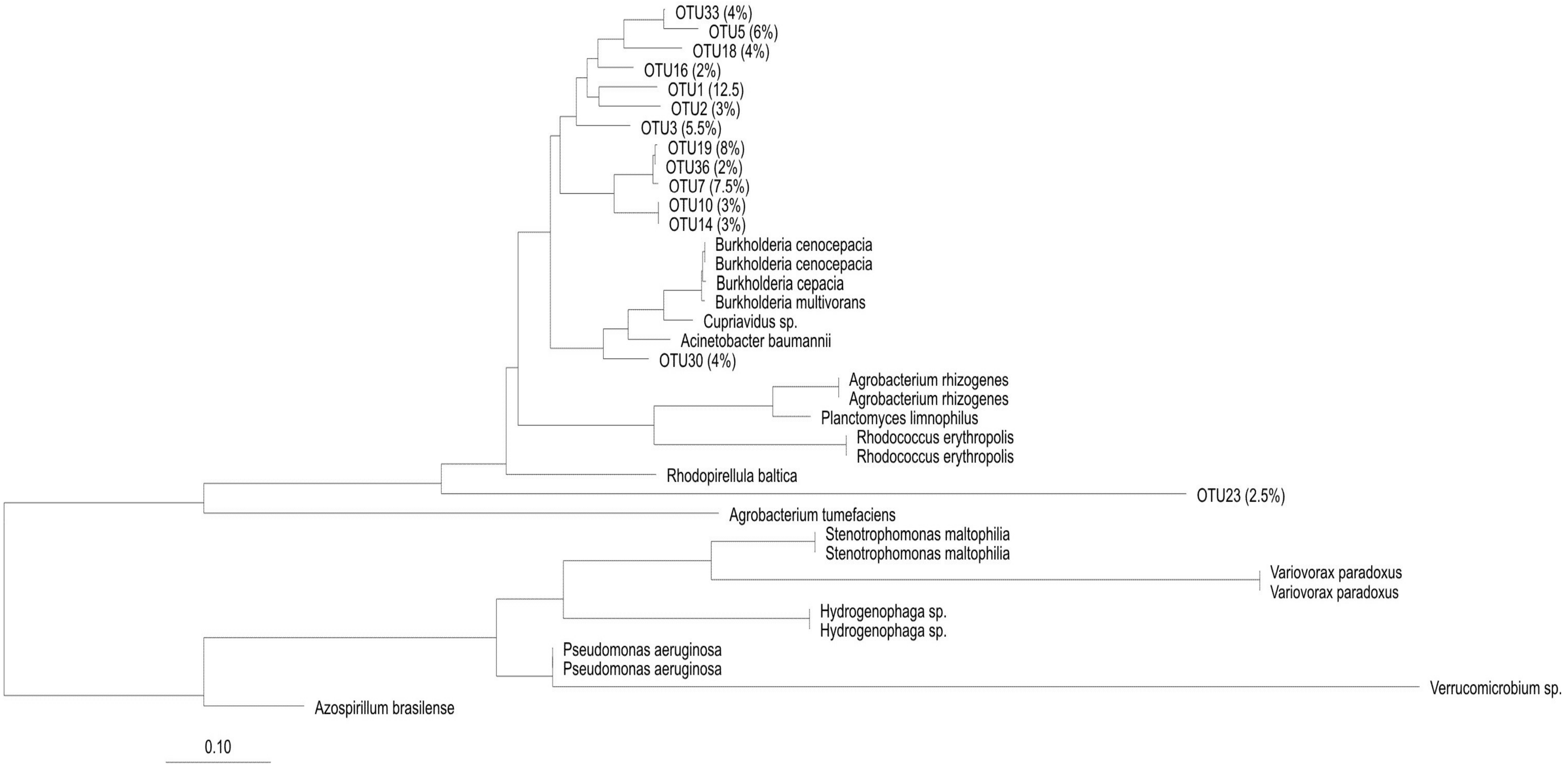


**Figure 3 – Randomised axelerated maximum likelihood tree of truncated AtsA protein sequences obtained from this study (Operational taxonomic units (OTUs)) and from described isolates in GenBank. The percentages (%) represent the overall abundance of the respective OTU in the *atsA* clone library.**

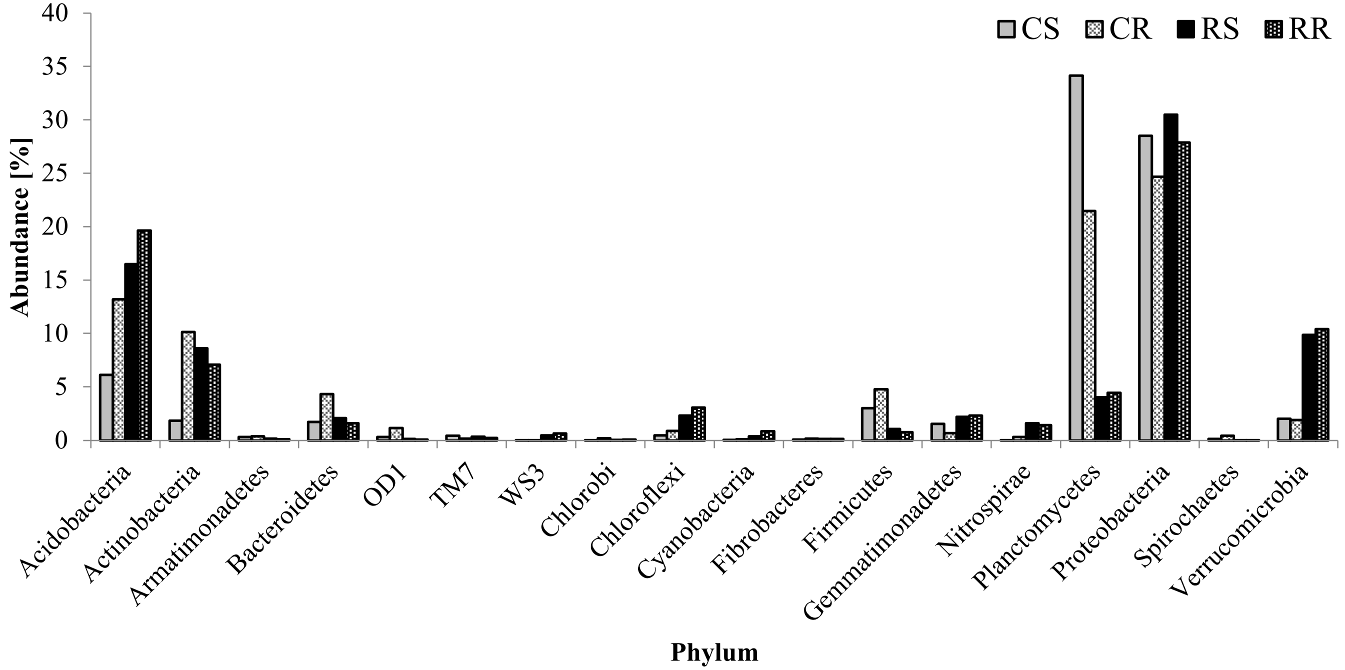


Supplementary Figure S9 – Abundance of sequences allocated to major bacterial phyla (cut-off 0.1%) after taxonomic analysis of 16S rRNA amplicons from hyphosphere cores (C) (grey) and rhizosphere roots (R) (black) of *Agrostis stolonifera* and static (S) = mycorrhizal (solid) and rotating (R) = severed mycorrhizal hyphae (pattern) treatments.

Supplementary information

#### *S1 - Determination of $^{34}$S uptake*

Determination of $^{34}$S uptake was achieved using Elemental Analysis - Isotope Ratio Mass Spectrometry (EA-IRMS) undertaken by Iso-Analytical (Chesire, UK). EA-IRMS is a technique used to determine the relative abundance of isotopes in a particular sample. Samples must be introduced to the mass spectrometer as pure gases and this was achieved via combustion. Tin capsules containing reference or sample material plus vanadium pentoxide catalyst were loaded into a furnace (1080 °C). The tin capsules flash combust in the presence of $O_2$ and the temperature was raised to 1700 °C. Following combustion, the gases were swept in a helium stream over combustion catalysts (tungstic oxide/zirconium oxide) and through a reduction stage of high purity copper wiring to produce $SO_2$, $N_2$, $CO_2$, and $H_2O$. A Nafion™ membrane was used to remove $H_2O$ and $SO_2$ was separated from $N_2$ and $CO_2$ on a packed GC column (32 °C).

The resultant pure $SO_2$ entered the ion source of the IRMS where it was ionised and accelerated. Separation of gas species based on mass was achieved in a magnetic field. Simultaneously, the mass-to-charge ration ($m/z$) of the different ion beams was measured on a Faraday cup universal collector array. Analysis was based on monitoring the $m/z$ at 48, 49 and 50 of $SO^+$ ions produced from $SO_2$ in the ion source to ascertain the relative abundance of $^{32}$S (48 $m/z$) to $^{34}$S (50 $m/z$). The total S content was calculated from the sum of the ion beams at $m/z$ 48, 49 and 50 which represent $^{32}$S, $^{33}$S and $^{34}$S, respectively. Both references and samples were converted to pure $SO_2$ and analysed using this method.

The reference material used for sulfur isotope analysis was IA-R061 (barium sulfate, $\delta^{34}S_{V\text{-}CDT}$ = +20.33‰). To calibrate and correct for the $^{18}$O contribution to the $SO^+$ ion beam IA-R061, IA-R025 (barium sulfate, $\delta^{34}S_{V\text{-}CDT}$ = +8.53‰) and IA-R026 (silver sulfide, $\delta^{34}S_{V\text{-}CDT}$ = +3.96‰) were used. Delta Units (δ) are expressed in molecules per thousand and are used to denote isotope ratios. For example, if $\delta^{34}S_{V\text{-}CDT}$ = 3.96‰ this means that the sample was analysed against a reference material and found to have 3.96 molecules per thousand more than V-CDT (Vienna-Canyon Diablo Troilite), an iron-sulfide meteorite, which is the accepted zero point for expression of $\delta^{34}S$.

*S2 - S K-edge X-ray absorption near edge spectroscopy*

Powdered samples were mounted on a sample holder for S K-edge XANES analysis. The monochromator of the beamline was operated in a step by step mode using a Si111 crystal. An ionisation chamber was used to measure the primary flux and a 4 element Si drift detector was used to measure the fluorescence signal. The X-ray energy was calibrated to elemental S at 2472 eV and scans were carried out from 2450 – 2520 eV in steps of 1 eV (2450-2468) and 0.5 eV (2468-2520) (10-20 s per step) in order to identify the different oxidation states of S. Recorded spectra were normalised after base-line subtraction and a linear combination fit was carried out using Athena software (Demeter package 0.9.16) (Solomon *et al.*, 2003).

*S3 - Percentage root colonisation*

A modified version of the grid line intersect method was used (McGonigle *et al.*, 1990) as follows. A representative population of roots (2 root fragments, 10 cm in length were picked from the top, middle, and bottom of the microcosms) and were cut into 1 cm segments. The roots were stained with 20 mL of 10% KOH (w/v) for 12 h. The segments were washed with $dH_2O$ and covered with 20 mL of alkaline $H_2O_2$ to bleach for 60 min. The bleaching solution was discarded and the roots were rinsed thoroughly with water. Roots were acidified in 20 mL of 0.1 M HCl solution for 12 h to ensure staining of intracellular fungal structures. The HCl solution was discarded and the roots were covered with 20 mL of lactoglycerol trypan blue stain (lactic acid: glycerol: $H_2O$ in a 1:1:1 ratio, with 0.05% (w/v) trypan blue) and incubated at 90 ºC for 45 min. The stained roots were then removed and covered in 20 mL of lactoglycerol destain (minus trypan blue) overnight prior to examination. The 1 cm root segments were examined one field of view at a time (x 1000 magnification). The field of view was moved in reference to a graticule inserted into the microscope eyepiece and the point of intersection was determined at the position of the graticule's vertical crosshair entering the root (Chapter 3, Section 3.2.2., Figure 17).

Once the point of intersection was noted, the field of view was moved completely through the root and the presence of (1) arbuscules (2) vesicles and (3) hyphae was noted as 'negative' (no fungal structures), 'arbuscules', 'vesicles', or 'hyphae only'. If the crosshair cut an arbuscule or vesicle, the respective category was increased by one and the total number of intersections was also increased by one. This was also the case for the 'hyphae only' category. In the case of both arbuscules and vesicles being recorded at an intersection, the individual categories were both incremented but the total number of intersections was increased only by one. AC and VC, respectively, were calculated by dividing their respective counts by the total number of intersections examined. HC was calculated as a proportion of the non-negative intersections.

*S4 - Minimal Media 2 with Toluenesulfonate or Lignosulfonate*

In this project, minimal media 2 with toluenesulfonate (MM2TS), lignosulfonate (MM2LS) and minimal media 2 without S (MM2SF) modified by Schmalenberger et al. 2008 were used to cultivate and compare growth rates of sulfonate mobilizing bacteria (Beil et al. 1995, Schmalenberger et al. 2008).

**10 x Tris stock N**

| Ingredient | Manufacturer | Product Code | g/L |
|---|---|---|---|
| Tris | Fisher Scientific | T/3710/60 | 15.138 |

| | | | |
|---|---|---|---|
| $NH_4Cl$ | Fisher Scientific | A/3920/53 | 10.7 |
| $MgCl_2$ (1 M) | Fisher Scientific | M/0600/53 | 5 ml |

**MMtris2 Medium**

| Ingredient | Manufacturer | Product Code | mL/L |
|---|---|---|---|
| Water | NA | NA | 825 |
| 10 x Tris stock N | Above | Above | 100 |
| Succinate (1 M) | Fisher Scientific | S/6490/48 | 5 |
| Glycerol (50 %) (v/v) | Fisher Scientific | G/0650/08 | 980 µL |
| Sodium /Potassium Chloride (1 M) | Fisher Scientific | BP366-500 | 20 |
| Potassium Phosphate (1 M) | Fisher Scientific | 215472500 | 10 |
| 200 x Trace elements | Section A6.3.1 | | 5 |
| *Toluene Sulfonate (100 mM, pH 7) **MM2TS** | Merck | 109613.0100 | 2.5 |
| *Lignosulfonate (100mM, pH 7) **MM2LS** | Aldrich | 471054 | 5.96 |
| $^{\beta}$Agar | Amresco | J637 | 6 |
| $^{\alpha}$Fructose (0.5 M) | VWR | 103637Y | 5.6 |

*Not included for S-free minimal media $^{\alpha}$added after autoclaving $^{\beta}$Solid media only

Trace Elements Stock

The trace element solution used was originally designed for cultivation of phototrophic sulfur bacteria (Pfennig and Lippert 1966).

**Trace Elements Stock**

| **Ingredient** | **mg/L** |
|---|---|
| $Na_2EDTA.2H_2O$ | 500 |
| $FeCl_2.4H_2O$ | 143 |
| $ZnCl_2$ | 4.7 |
| $MnCl_2.4H_2O$ | 3 |
| $H_3BO_3$ | 30 |
| $CoCl_2.2H_2O$ | 20 |
| $CuCl_22H_2O$ | 1 |
| $NiCl_2.6H_2O$ | 2 |
| $Na_2MoO_{4.}2H_2O$ | 3 |
| $CaCl_2.2H_2O$ | 100 |

The MM2TS, MM2LS or MM2SF solution was autoclaved for 15 min at 121° C for sterilisation. Subsequently, 5.6 ml/L of 0.5 M sterile fructose (pH 7.2) was added to the mixture. The fructose is added after autoclaving to prevent decomposition of the compound. Additionally, for solid media, 4.8 g/L of agarose standard was added prior to autoclaving.

*S5 – PCR-DGGE*

Bacterial 16S rRNA gene amplification was carried out with this DNA using the primer pair GC-341F/518R (supplementary Table S1) targeting the V3 region for DGGE (Muyzer *et al.*, 1993). The final concentration per 25 µL reaction was 1 X buffer (2 mM $MgCl_2$), 0.2 mM dNTP mix, 0.4 µmol of each primer, and 0.5 U of DreamTaq polymerase (Fermentas, Waltman, MA). A touchdown PCR protocol was used with the following cycling conditions: initial denaturation of 94 °C for 5 min, 20 cycles of 94 °C denaturation (45 s), 65-55 °C touchdown (45 s), 72 °C extension (45 s), plus 18 further cycles with an annealing temperature at 55 °C. Final extension was carried out at 72 °C for 5 min. DGGE was carried out on 200 x 200 x 1 mm gels in a TV400 DGGE apparatus (Scie-Plas, Cambridge, UK). Gels of 10% (w/v) acrylamide/bisacrylamide were prepared and run using a linear 30-60% gradient in 1 X TAE buffer (60 °C) for 16.5 h at 63 V (Fox *et al.*, 2014). After completion, gels were stained with SYBR Gold (1:10,000 diluted) (Invitrogen, Carlsbad, CA) for 30 min and the image captured on a Syngene G:Box (Cambridge, UK).

AM fingerprinting was achieved using the AM specific primer AM1 (Helgason *et al.*, 1998) alongside the universal eukaryotic primer NS31 (Simon *et al.*, 1992) (Table 11) targeting 18S rRNA. The PCR were undertaken in 25 µL reactions with 1 X buffer (2 mM $MgCl_2$), 1 M betaine, 0.2 mM dNTP mix, 0.4 µmol of each primer and 0.5 U of DreamTaq polymerase (Fisher Scientific, Waltham, MA). The PCR was carried out under the following conditions: initial denaturation of 94 °C for 5 min, 30 cycles of 94 °C denaturation (30 s), 58 °C annealing (60 s), and 72 °C extension (90 s). Final extension was carried out at 72 °C for 5 min. A tenfold dilution of the PCR product was undertaken and used as template for a nested PCR using the primer set Glo1 (Kowalchuk *et al.*, 2002) and NS31-GC (Cornejo *et al.*, 2004) (Table 11). The nested PCR was carried out under the following conditions: initial denaturation of 94 °C for 5 min, 20 cycles of 94 °C denaturation (45 s), 58-48 °C touchdown (45 s), 72 °C extension (45 s), plus 15 further cycles with an annealing temperature at 48 °C. Final extension was carried out at 72 °C for 5 min. DGGE was run as before with a gradient of 35-55%.

Fungal DNA fingerprinting was undertaken using the fungal specific primer ITS-1F (Gardes & Bruns, 1993) and ITS-4 (White *et al.*, 1990) (Table 11). This product was tenfold diluted and used as template in a nested PCR using an ITS-1FGC primer with a 40 base GC clamp to the 5' end of the primer (Bougoure & Cairney, 2005) and ITS-2 reverse primer (White *et al.*, 1990) (Table 11). Both PCRs were undertaken in 25 µL reactions with 1 X buffer (2 mM $MgCl_2$), 1 M betaine, 0.2 mM dNTP mix, 0.4 µmol of each primer and 0.5 U of DreamTaq polymerase (Fisher Scientific, Waltham, MA). For both PCR reactions, amplification was performed with an initial denaturation of 94 °C for 5 min, 40 cycles with the first 20 cycles at 95 °C denaturation (45 s), 60 °C annealing (45 s), and 72 °C extension (45 s). Cycles 21–40 used the same parameters with annealing temperature of 50 °C (Gardes & Bruns, 1993). Final extension was carried out at 72 °C for 5 min. DGGE was run as before with a gradient of 35-65%.

Table S1

**Primers used in this study**

| ID | Sequence | Target | Reference |
|---|---|---|---|
| 27F | 5'-AGAGTTTGATCMTGGCTCAG-3' | 16S Rrna | Lane 1991 |
| 1492R | 5'-GGTTACCTTGTTACGACTT-3' | 16S Rrna | Lane 1991 |
| AsfAF2 | 5'-TACATGCGSCTGATGCGCA-3' | *asfA* gene | Schmalenberger and Kertesz 2007 |
| AsfBtoA | 5'-ASCTCGCACATGAAGCAGG-3' | *asfA* gene | Schmalenberger and Kertesz 2007 |
| AtsA-F1 | 5'-TIGCIGAYGAYITSGGITWYTCTGA-3' | *atsA* gene | M. Kertesz and A. Houlden |
| AtsA-R1 | 5'-TCSGSICCRTTGTCSGACATGAA-3' | *atsA* gene | M. Kertesz and A. Houlden |
| 341F-GC | 5'CGCCCGCCGCGCGCGGCGGGCGGGGCGGGGGCACGGGGGGCCTCGGG ACCGAGCAG-3' | 16S Rrna | Muyzer et al. 1993 |
| 518R | 5'-ATTACCGCGGCTGCTGG-3' | 16S rRNA | Muyzer et al. 1993 |
| Glo1 | 5'-GCCTGCTTTAAACACTCTA-3' | 18S rRNA | Kowalchuk et al. 2002 |
| NS31-GC | 5'CGCCCGGGGCGCGCCCCGGGCGGGGCGGGGGCACGGGGGTTGGAGGG AGGGCAAGTCTGGTGCC-3' | 18S rRNA | Cornejo et al. 2004 |
| ITS1F | 5'-CTTGGTCTTTAGAGGAAGTAA-3' | ITS fungal | Gardes and Bruns 1993 |
| ITS4 | 5'-TCCTCCGCTTATTGATATGC-3' | ITS fungal | White et al. 1990 |
| ITS1F-GC | 5'CGCCCGCCGCGCGCGGCGGGCGGGGCGGGGGCGCGGGGGCCTTGGTC TTTAGAGGAAGTAA-3' | ITS fungal | Bougoure and Cairney 2005 |
| ITS2 | 5'-GCTGCGTTCTTCATCGATGC-3' | ITS fungal | White et al. 1990 |

| | | | |
|---|---|---|---|
| AM1 | 5’-GTTTCCCGTAAGGCGCCGAA-3’ | 18S AM fungal | Helgason et al. 1998) |
| NS31 | 5’-TTGGAGGGCAAGTCTGGTGCC-3’ | 18S rRNA | Simon et al. 1992 |
| asfAF1all | 5’-YTSTCVGGCATGGAGTTYT-3’ | *asfA* gene | Gahan and Schmalenberger 2015 |
| 16SF | 5’TCGTCGGCAGCGTCAGATGTGTATAAGAGACAGCCTACGGGNGGCWGCAG 3’ | 16S (V3-V4 region) | Klindworth et al. 2012 |
| 16SR | 5’CTCTCGTGGGCTCGGAGATGTGTATAAGAGACGAGACTACHVGGGTATCTAATCC -3’ | 16S (V3-V4 region) | Klindworth et al. 2012 |
